# Evaluating Physiological Indicators in Detecting Deception and Truthfulness Using the Comparison Question Test

**DOI:** 10.1101/2024.09.01.609623

**Authors:** A M Shahruj Rashid, Bryan Carmichael, Charlize Su, Keming Shi, Keefe Lim, Senthil Kumar Poorvika, Ngok Jeun Wan, Eshaan Govil, Dennis Yap

## Abstract

Despite significant advancements in deception detection, traditional methods often fall short in real-world applications. This study addresses these limitations by evaluating the effectiveness of various physiological measures — pupil response, electrodermal activity (EDA), heart rate (HR), and facial temperature changes — in predicting deception using the Comparison Question Test (CQT). It also fills a critical research gap by validating these methods within an Asian context. Employing a between-subjects design, data was collected from a diverse sample of 118 participants from Singapore, including Chinese, Indian, and Malay individuals. The research aims to identify which physiological indicators, in combination, offer the most robust predictions of deceptive behavior. Key innovations include the adaptation of the CQT with a modified directed lie paradigm and an expanded sample size to assess the relative importance of each physiological measure. The study’s findings reveal that pupil response is the most significant predictor of deception, with EDA enhancing the model’s explanatory power. HR, while relevant, adds limited value when combined with pupil response and EDA, and facial temperature changes were statistically non-significant. The study highlights the need for further research into the interactions among physiological measures and their application in varied contexts. This research contributes valuable insights into improving deception detection methodologies and sets the stage for future investigations that could incorporate additional physiological indicators and explore real-world applications.

## 1. Introduction

Reliable deception detection is crucial in uncovering crimes,establishing credibility, and has significant applications in law enforcement, security, counterterrorism, and job applicant screening. In places like Belgium, Japan, India, Taiwan and Indonesia, data from deception detection devices are even admissible as legal evidence. Traditionally, deception detection has relied heavily on human interpretation of polygraph results or behavioral tests, which can achieve up to 90% accuracy rates in controlled environments (Nelson, 2015).

However, outside the domain of polygraph experts, the efficacy of deception detection diminishes significantly. Studies consistently demonstrate that when individuals attempt to identify deception without specialized tools, their accuracy hovers just above chance levels, averaging around 54% (Bond and DePaulo, 2006; Mann et al., 2004). More specifically, when individuals are tasked with distinguishing between truth and deception solely by relying on their own perceptions, they accurately identify lies as false only 47% of the time and correctly recognize truth as non-deceptive in approximately 61% of cases (Vicianova, 2015). The stark gap between potential and actual accuracy highlights a critical shortfall in human lie detection capabilities, fuelling a surge in interest surrounding technology that can assist in the process of lie detection, including next-generation modalities such as infrared (IR) and thermal sensors.

While automated deception detection technologies have shown promise in some studies, a significant gap remains in the literature — particularly regarding the validation of these technologies within Asian populations (O’Shea et al., 2018; Bittle, 2020). Addressing this gap is crucial for ensuring that these technologies are universally applicable and effective across diverse cultural contexts.

The current study aims to bridge this gap by adopting a between-subjects design to collect data encompassing interconnected physiological aspects to determine which parameters, in combination, best predict deception based on known ground truths. It aims to assess the effectiveness of our adaptation of the established Comparison Question Test (Nelson and Handler, 2007) testing techniques in detecting deception and ultimately, enhancing deception detection technologies. The central objectives of this study are to test combinations of features for deception detection, validate these methods in an Asian population, and replicate prior research with significant variations. These variations include the use of a modified directed lie paradigm, a larger sample size, and a Singaporean sample composed mainly of ethnically Chinese, Indians, and Malays. Given the increasing evidence that broadly multimodal approaches to deception detection may be superior compared to approaches that rely on a limited set of indicators (Pérez-Rosas et al., 2014), this study is well-positioned to contribute to valuable insights. To ensure transparency and rigor in our research design, our study was pre-registered on AsPredicted (#151764).

### 1.1 Literature Review

#### 1.1.1 Eye Features (Pupil Diameter)

Pupil dilation and constriction are physiological responses of the autonomic nervous system (ANS) that reflect cognitive and emotional states. Research has demonstrated that cognitive and emotional manipulations involve ANS responses, which are reflected in changes in pupil size (Oliva and Anikin, 2018). For instance, pupil dilation can be influenced by factors such as deception, where the cognitive load and emotional stress associated with lying lead to observable changes in pupil size (Webb et al., 2009). The modulation of pupil size is governed by the sphincter and dilator muscles, which, in turn, are under the control of the ANS. Specifically, sympathetic activities, a component of the ANS, escalate in response to stressful situations, culminating in an increase in pupil dilation (Kupcová, 2017). Such ANS-driven processes, including pupil dilation, occur involuntarily.

An augmentation in pupil dilation typically signifies heightened cognitive load (Szulewski et al., 2014). Deception inherently involves an escalation in cognitive demands (Vrij et al., 2010). When individuals engage in deception, they often intricately construct a narrative that adheres to truth while also appearing logical and believable. As a result, the sustained suppression of the truth throughout the narrative significantly elevates cognitive demands. Therefore, it is plausible that pupil dilation may be an indicative marker of deceptive conduct.

In Dionisio et al.’s research conducted in 2001, participants were tasked with responding to questions, alternating between providing truthful and deceptive answers. Notably, when participants were fabricating responses, their pupils exhibited a significantly greater dilation compared to instances when they were recounting truths from their memory. These findings suggest a potential association between increased pupil size and engagement in deceptive behaviors. Additionally, individuals attempting deception exhibited significantly larger pupillary responses when confronted with relevant questions compared to those who answered truthfully (Berrien and Huntington, 1943). Other contributions in the literature, including studies by Leal and Vrij (2008), Webb et al. (2009), Walczyk et al. (2012), and Vrij et al. (2015) have further proposed a direct connection between the act of crafting lies and observable oculomotor patterns such as blinking, fixations, saccades, and pupil dilation. Eye features, especially pupil dilation, are best recorded using an eye tracker or IR camera together with IR lighting.

In a study by Kuhlman et al. (2011), the potential of pupil diameter (PD) changes as indicators of deception was investigated. This research builds on the premise that changes in PD are associated with cognitive load (Beatty, 1982) and emotional arousal (Bradley and Janisse, 1979; Janisse and Bradley, 1980), making them viable markers for detecting deceit. The study involved 112 subjects, divided into innocent and guilty groups, who answered a series of statements while their PD was recorded using an eye tracker. The results indicated significant habituation in PD responses over repeated sessions, with the mean slope of regression lines for pupil response amplitude decreasing across the first three repetitions. Despite this habituation, the diagnostic validity of using PD changes for deception detection remained stable, suggesting that while emotional arousal may decrease, the cognitive effort required for deception continues to affect PD. This study supports previous findings by Vrij et al. (2011), highlighting the relevance of PD in lie detection and emphasizing the need for further research into the effects of task instructions on diagnostic validity in the presence of habituation.

Overall, pupil dilation involves both sympathetic and parasympathetic inputs and is highly sensitive to novelty, motivational significance, and the intensity of stimuli (Nieuwenhuis et al., 2011). The consistent findings across various studies indicate that pupil size changes, particularly dilation, serve as reliable markers of cognitive load and emotional arousal. These physiological responses, governed by the ANS, reflect the underlying cognitive and emotional states associated with deception. The elevated cognitive demands of lying lead to notable increases in pupil size, distinguishing deceptive behavior from truthful responses. The research underscores the potential of pupil dilation as an effective indicator in lie detection, highlighting its significance in understanding and measuring the cognitive and emotional processes involved in deception.

#### 1.1.2 Heart Rate

Heart rate regulation is primarily governed by the ANS, which comprises the sympathetic and parasympathetic branches. The sympathetic nervous system (SNS) accelerates heart rate through the release of norepinephrine, which binds to beta-adrenergic receptors on the heart, enhancing the rate and force of contractions. In contrast, the parasympathetic nervous system (PNS), via the vagus nerve, exerts a braking effect on the heart by releasing acetylcholine, which binds to muscarinic receptors, slowing the heart rate. Following an initial acceleration of heart rate driven by sympathetic activation, the subsequent reduction in heart rate can occur as parasympathetic activity increases and sympathetic activity decreases.

Factors such as respiratory behavior and baroreflex activation also modulate heart rate changes, where the baroreflex mechanism helps in maintaining blood pressure stability by adjusting heart rate in response to changes in blood vessel stretch. This dynamic balance between the SNS and PNS allows for rapid adjustments in heart rate in response to physiological demands, such as during stress or relaxation.

Additionally, the interaction between these systems is influenced by central control mechanisms located in the brainstem, particularly within the medulla oblongata, which integrates sensory input and modulates autonomic output to maintain cardiovascular homeostasis (Nieuwenhuis et al., 2011). Deceptive behavior often triggers physiological arousal reminiscent of the “fight or flight” response. This heightened arousal can manifest as an acceleration of heart rate, as demonstrated in studies such as the one conducted by Verschuere et al. (2004). Deception also frequently elicits emotional states such as anxiety, fear, or guilt, which can further influence heart rate, with increased heart rate often associated with heightened emotional reactions (Honts et al., 1996).

Podlesny and Raskin’s (1977) seminal work provided crucial insights into the relationship between guilt and heart rate in the context of deception detection. Their meta-analysis revealed that guilty subjects consistently exhibited a marked reduction in heart rate following their responses to pertinent questions. This decrease is likely due to the activation of the PNS. Lying often induces stress and anxiety, triggering the SNS’ “fight or flight” response. To counterbalance this, the PNS is activated, helping to bring the body back to a state of calm and homeostasis. Additionally, lying can evoke strong emotions such as guilt, fear, and anxiety. The activation of the PNS as part of the emotional recovery process helps individuals calm down and manage these negative emotions, further aiding in the restoration of physiological balance.

Klein Selle et al. (2015) delved into the nuances of heart rate responses in deception scenarios. The orienting response (OR) and arousal inhibition (AI) theories offer distinct explanations for the physiological responses observed during the Concealed Information Test (CIT). The OR theory posits that the presentation of novel or significant stimuli, such as crime-related details, triggers an immediate orienting response characterized by SNS activation. This activation leads to increased skin conductance and an initial heart rate acceleration followed by a brief deceleration, mediated by transient PNS activation.

In contrast, the AI theory emphasizes the sustained efforts of individuals to inhibit their physiological arousal in order to conceal information. This ongoing process predominantly involves sustained SNS activation to manage arousal and continuous PNS activation to maintain prolonged heart rate deceleration. Thus, while the OR theory explains physiological responses as short-lived reactions to significance and novelty, AI theory accounts for more enduring responses driven by the need to suppress arousal, highlighting the distinct roles of SNS and PNS in these mechanisms.

In their study, participants mentally immersed themselves in assigned roles (as either suspects or witnesses to a burglary), suspects showed a more significant decline in heart rate compared to witnesses, emphasizing the role of emotional responses in heart rate modulation during deception. Klein Selle et al. (2015) argued that heart rate deceleration is primarily driven by AI rather than OR. While suspects showed significant heart rate deceleration in response to relevant items, witnesses did not exhibit this pattern. Together, these studies illuminate the intricate interplay between deception, emotional states, and heart rate dynamics, with the groundwork laid by Podlesny and Raskin (1977) continuing to shape our understanding of this phenomenon.

#### 1.1.3 Thermal Imaging

Utilizing thermal imaging for deception detection involves employing a thermal camera to gauge the temperature of facial skin as an indicator of potential deception. Research has shown that deceptive behavior can be correlated with significantly higher facial temperatures, particularly in the periorbital regions (Park et al., 2013). Moliné et al.’s 2018 study investigates the thermographic correlates of lying, planning, and anxiety. Using high-resolution IR thermography, the researchers measured facial temperature changes in participants subjected to tasks designed to induce these specific cognitive-emotional states. The study focused on the temperature variations in the nose and forehead regions, which are highly sensitive to ANS activity.

The study’s most prominent finding is the “Pinocchio effect,” wherein lying leads to a significant decrease in nose temperature. This phenomenon is explained by the ANS’ response to the psychological stress and cognitive load associated with deception, corresponding with increased activity and blood flow in the brain regions involved in deceptive behavior. This consistent physiological response to lying suggests that thermography could be a valuable tool for non-invasive lie detection. In addition to deception, the research explored the thermographic effects of cognitive load associated with planning. Participants engaged in tasks requiring substantial mental effort exhibited an increase in forehead temperature. This thermographic change is attributed to increased neural activity and cerebral blood flow in the prefrontal cortex, a region essential for complex cognitive functions.

Conversely, anxiety-induced tasks resulted in variable facial temperature changes, reflecting the heterogeneity in individual physiological responses to stress. The study underscores the complexity of anxiety’s thermographic manifestations, influenced by both task-specific and individual differences. These findings provide a compelling rationale for developing computer vision-based lie detection systems that utilize thermography. By capturing real-time facial temperature changes, such systems could offer a non-invasive and immediate assessment of an individual’s truthfulness.

#### 1.1.4 Skin Response

The galvanic skin response (GSR), alternatively known as EDA or skin conductance response, is a psychophysiological phenomenon reflected in electrical conductance changes as sweat glands are activated. This physiological reaction is intricately tied to the modulation of sweat gland activity by the ANS. Notably, skin responses are exclusively sympathetic; there is no parasympathetic innervation of sweat glands. When an individual engages in deception or falsehood, discernible shifts in their physiology may manifest, characterized by an augmentation in perspiration. This perspiration, in turn, exerts an influence on skin conductivity. The monitoring of GSR, therefore, becomes a method of capturing these nuanced changes. The underlying premise for employing GSR as an indicator of deception lies in the notion that deception can elicit heightened emotional arousal or stress, subsequently precipitating alterations in skin conductance (Vincent and Furedy, 1992). The meticulous measurement of these changes through GSR serves as a potential means of discerning deceptive states. GSR is a stimulus-locked EDA, typically measured using electrodes placed on the hands, that reflects increased sympathetic activity (Dawson et al., 2017).

Geldreich E.W. found that skin conductance was significantly increased during subjects’ deceptive moments compared to truthful ones, attributing this rise to autonomic disruptions and the patterned association of such disturbances with deceptive behavior. In polygraph examination techniques most frequently used in recent times, for instance, The Utah Zone Comparison Technique (UZCT), in the numerical assessment of the recordings, the diagnostic value of GSR is assessed much higher than respiratory measures (Widacki, 2015).

#### 1.1.5 Respiration

Respiratory line length (RLL) is a quantitative measure used by polygraphers to assess the complexity and variability of respiration (RSP), calculated by summing the vertical distances between consecutive points in the respiration signal over a specific time interval (Timm, 1982). A decrease in RLL has been linked to heightened arousal during deception, as individuals may unconsciously alter their breathing patterns, leading to shallower and more irregular breaths (Matsuda, 2011). Changes in these measures can signal stress responses, such as suppression or reduction in breathing cycles, often relevant in detecting deceptive behavior. Modern approaches address biases in traditional respiration measurements by using weighted averages to adjust for variability in the inclusion of respiratory cycle segments within analysis intervals, leading to more accurate assessments (Matsuda, 2011). Respiration is considered one of the most easily manipulated physiological signals in traditional polygraph examinations. Since respiration is a consciously controllable process, individuals can deliberately alter their breathing patterns to attempt to deceive polygraph results (Honts et al., 1996).. Techniques such as adjusting the depth and rate of breathing can interfere with the detection of emotional and physiological responses typically associated with deception. For example, a subject might try to mask signs of stress by breathing deeply or maintaining a regular breathing pattern, thereby reducing the polygraph’s effectiveness in detecting changes related to stress or arousal. Moreover, respiration is closely linked to cardiac responses due to the interplay between the respiratory and cardiovascular systems, a phenomenon known as respiratory sinus arrhythmia (RSA). RSA refers to the natural fluctuation in heart rate that occurs during the respiratory cycle—heart rate increases during inhalation and decreases during exhalation. This connection implies that manipulating respiration can also influence heart rate measurements.

## 2. Methodology

### 2.1 Ethical Considerations

This study was conducted under the ethical oversight of the Parkway Independent Ethics Committee (PIEC), with IRB clearance granted under reference number PIEC/2022/062. The study, titled “Deception Detection with an Automated Interviewing System,” received approval for amendments on 10 October 2023, and was conducted in compliance with ICH GCP standards and applicable laws and regulations. This study is designed to minimize risks to participants, ensuring their well-being throughout the research process. While the study does not present inherent risks beyond those typically encountered in daily life, it involves elements of deception that may elicit emotional responses or discomfort in some individuals. To address these potential concerns, participants were fully informed of the nature of the study, including the use of deception, in the informed consent process. They were made aware of their right to withdraw from the study at any time without penalty. The informed consent document emphasized the importance of careful consideration before participation, allowing individuals to make an informed decision about their involvement.

### 2.2 Test Materials

The general paradigm of the Utah Directed-Lie Test (DLT), in which the participant is instructed to answer certain questions falsely (Raskin and Honts, 2002) was utilized. The DLT is a subtype of the Comparison Question Test (CQT), an “empirically consistent and unified approach to polygraphy” (Nelson and Handler, 2007). It is the deception detection questioning technique that is most widely used. We adapted and modified the Utah DLT (Raskin and Honts, 2002) for our Mock Crime, especially in the handling of the Directed Lie questions which we have replaced with the Moral Directed Lie Test (MDLT) questions. The DLT is designed to increase cognitive load, which is often higher in individuals being deceptive. By comparing the cognitive load induced by Directed Lie questions with that from relevant questions (Table 1), we can assess whether a participant is being truthful or deceptive. However, preliminary trials revealed that the current set of Directed Lie questions, which focused on participants’ life experiences, led to confusion. This confusion resulted in responses that were either subjective or truthful, compromising the experiment’s outcomes. To address this, we needed to design questions that were less likely to elicit subjective answers.

**Table 1:**
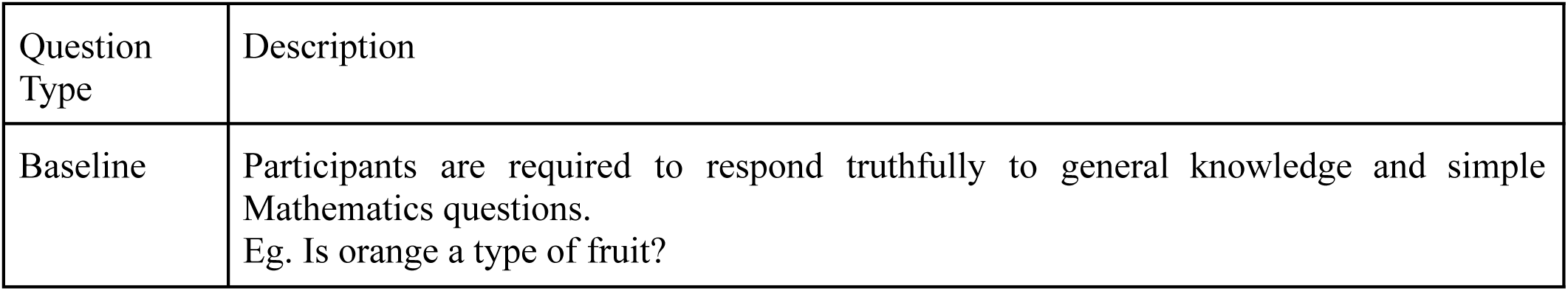
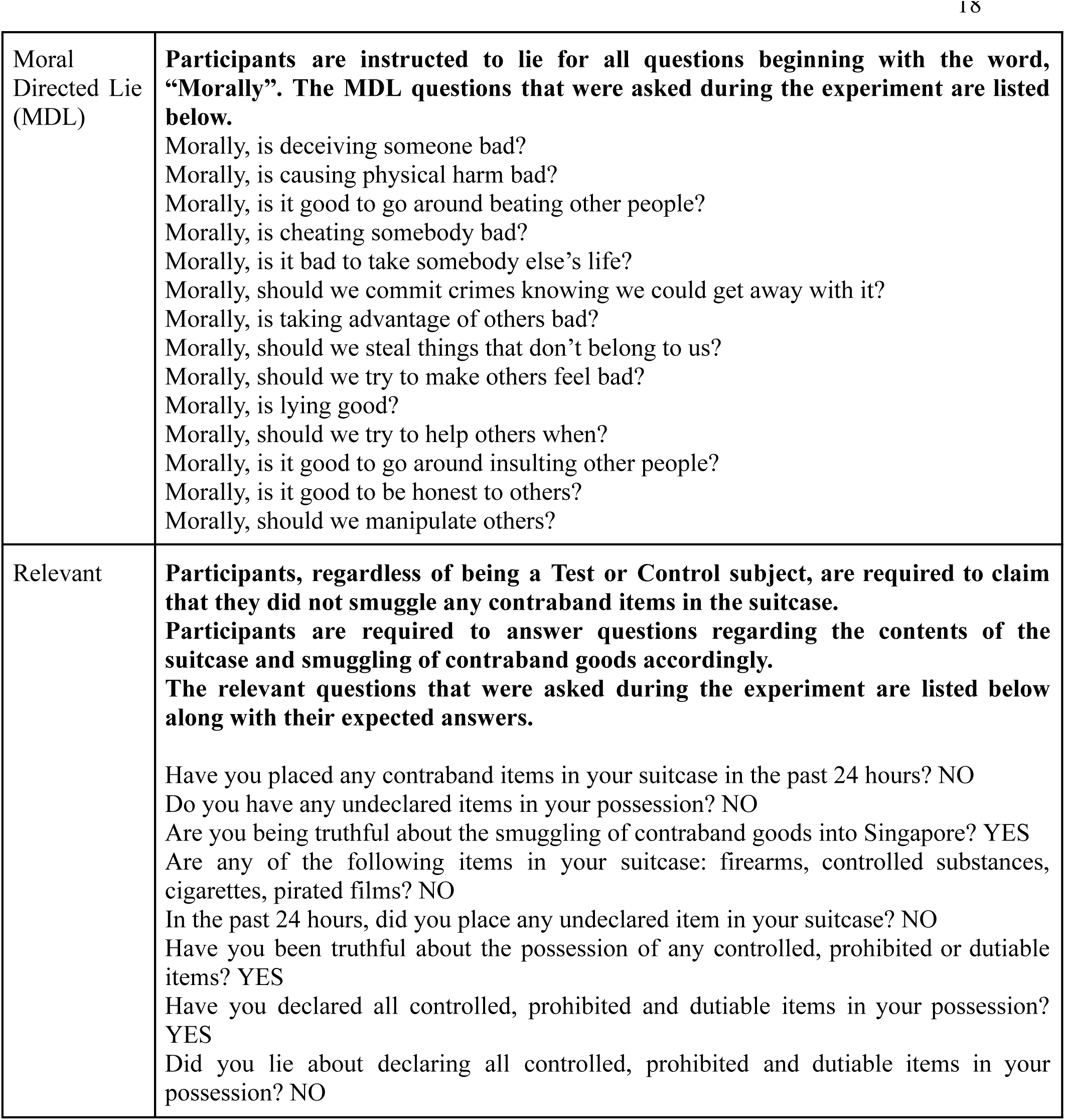
Question Types.

The Moral Directed Lie Test (MDLT) was inspired by the need for a counterpart to the traditional Directed Lie Test (DLT) that would be more universally applicable and less vulnerable to variability in participants’ personal histories. When individuals are under intense cognitive load, their decision-making shifts from intuitive to more deliberate, rational processing (Kahneman, 2013). Prior research by Kvaran et al. (2013) demonstrated that moral dilemmas—such as the trolley problem (Foot, 1967)—are effective at engaging rational thinking. This connection is directly relevant to deception, as lying requires higher-order cognitive functions, placing individuals in a state of elevated cognitive load. Building on this insight, the MDLT replaces life-experience-based questions with morality-based questions, aiming to provide a more consistent and interpretable baseline that reliably induces rational thinking across participants.

This design also addresses two practical limitations encountered in earlier work. In previous in-house experiments using the standard directed lie paradigm (n = 74; 37 test, 37 control), we successfully replicated the expected pupil-dilation differences across conditions. The outcome of the previous in house experiment for pupil dilation is shown in Table 7.

However, participants commonly reported confusion about the phrasing and intention of traditional DLT questions, which often referenced autobiographical events that did not apply equally across individuals. This introduced unnecessary variability and occasional non-compliance. These challenges motivated us to explore whether changing the control-question type—while preserving the core “instructed lie with known ground truth” structure—would reduce confusion without compromising the cognitive-load manipulation. The MDLT represents our first systematic attempt to evaluate how altering the question class influences participants’ cognitive state and, ultimately, psychophysiological responses.

A key conceptual distinction in the MDLT is that participants are not asked to lie about an objective fact from their personal life; instead, they are asked to lie about what they believe to be true in a moral context. Unlike the DLT, where the cognitive load partly arises from the complexity or autobiographical nature of the question, morality-based questions place participants in a state of high cognitive load regardless of their personal experiences. We hypothesize that this offers a more stable and less confounded method for generating a high-load comparison condition.

In the present study, the Mock Crime experiments used three question types: Baseline questions, Moral Directed Lie (MDL) questions, and Relevant questions. Table 1 describes these categories. Participants responded to 42 questions organized into triplets. Each triplet began with a Baseline question, followed by either a Relevant and then an MDL question, or the reverse ordering; the second and third positions were randomized to mitigate order effects. This sequencing is shown in Figure 1. A 15-second interval separated each question, and the full test lasted approximately 10–15 minutes. After completing the test, participants performed an N-back task progressing from 1-back to 3-back. Although collected to inform future work, these N-back data were not analyzed or included in the current study.

**Figure 1.**
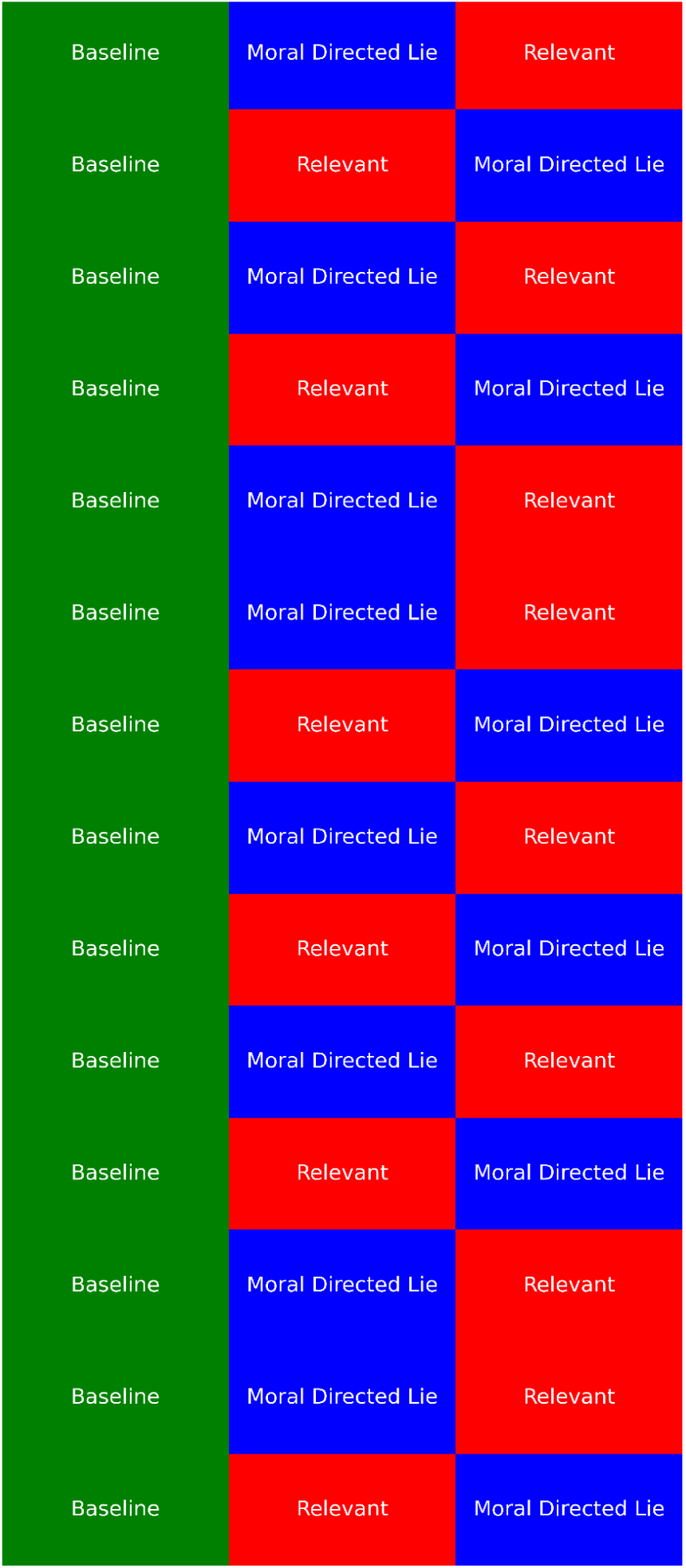
Pseudorandom Presentation of Question Types in Batches of 3 (Left to Right, Top to Bottom)

The data supporting the findings of this study are proprietary and belong to AI Seer Pte. Ltd. Due to confidentiality agreements and commercial sensitivities, the raw dataset cannot be publicly shared. However, a processed version of the data is available at <u>Harvard Dataverse</u>.

### 2.3 Design and Procedure

The ‘Mock Crime’ follows the structure of our adapted MDL and comprises 42 questions in which the participant is instructed to answer certain questions falsely. Participants were engaged in a role-playing scenario as individuals traveling to Singapore, who were randomly chosen for a routine customs inspection. Two experimenters played the roles of a narrator and a customs inspector. As shown in Table 2, the narrator explained the participant’s role by reading two predefined scripts, one for Test subjects and one for Control subjects. Half of the participants were randomly assigned as Test subjects, while the remaining half were Control subjects. Test subjects were instructed to choose from three envelopes containing various contraband goods, including drugs and narcotic substances. They were then given time to pack the chosen contraband item in the suitcase provided, trying to conceal it. Control subjects were also given time to pack the suitcase.

**Table 2.**
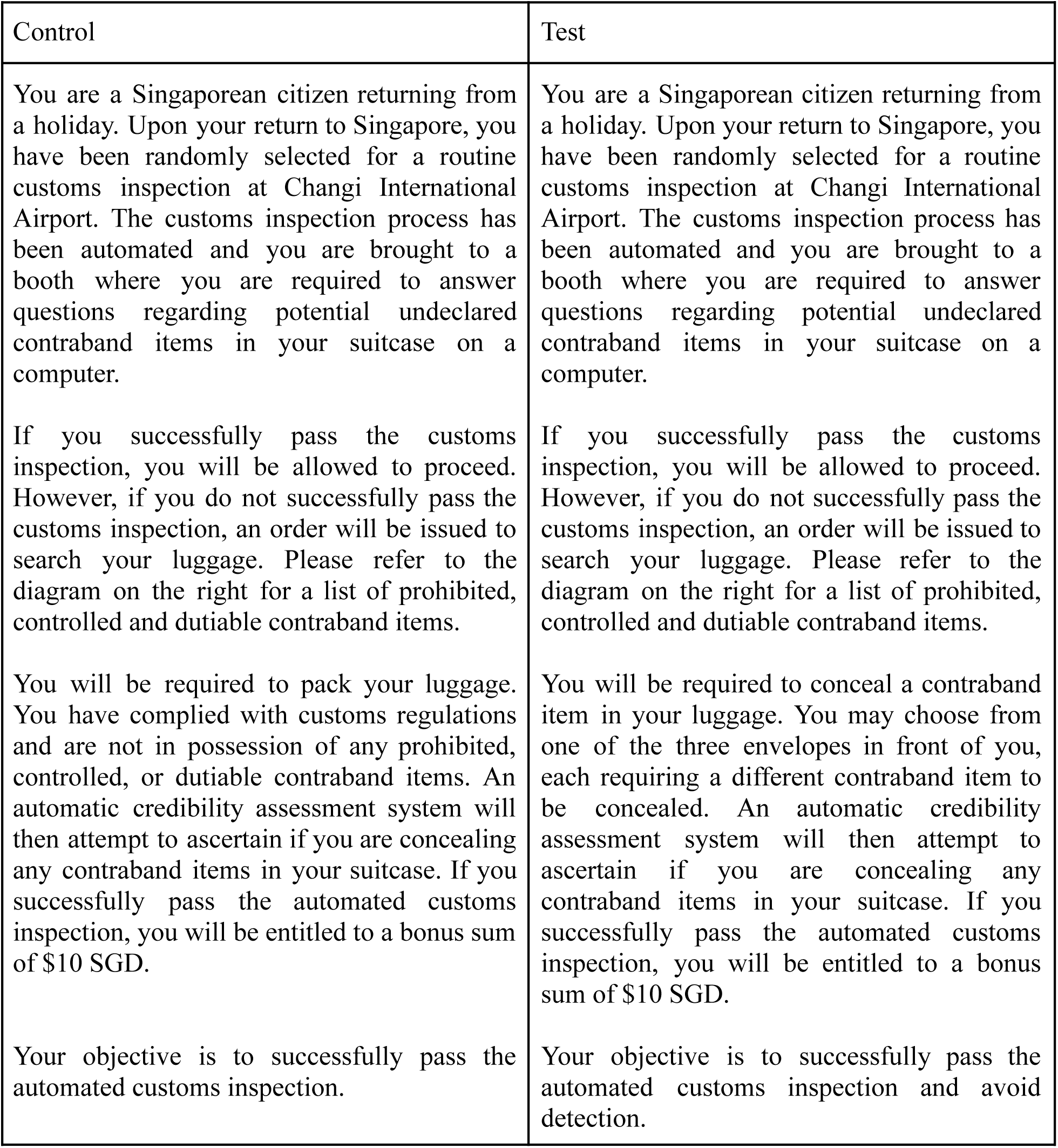
Predefined Scripts Read Out to Participants by the Narrator.

Participants were instructed to enter another room where the automated interview process would take place. The experimenter playing the role of the customs inspector will read out the following script:

You have been randomly selected for a routine customs inspection. This is in line with our efforts to curb the smuggling of contraband goods into Singapore. Before we begin, I would just like to remind you that contraband goods include items such as cigarettes, liquor, and drugs or other narcotic substances. Please refer to the diagram on the wall for a list of items that are prohibited, controlled and dutiable. Failure to declare the possession of such items will result in fines and/or prosecution. Should you have any of these items in your possession right now, please declare them. So, do you have any undeclared goods as listed previously in your possession?

Participants were outfitted with EDA, ECG, RSP, and PPG sensors to monitor physiological responses throughout the experiment. Thermal and RGB video data were collected. Their gaze was continuously tracked using an eye tracker. Once the calibration of all systems was completed, the experiment proceeded without further interaction from the experimenters. Three types of questions—baseline, MDL, and relevant—were presented in a randomized sequence, ensuring that no two questions of the same type appeared consecutively. Participants were instructed to respond with “YES” or “NO” to each question after it was read aloud by an automated voice. Responses were made using a mouse, with a left-click indicating “YES” and a right-click indicating “NO.”

### 2.4 Apparatus

Pupil diameter and gaze data were collected using the Tobii 5L eye tracker, operating at 120Hz. Respiration rates (RSP) and electrocardiography (ECG) signals, Photoplethysmography (PPG) and EDA were concurrently measured using the MP160 BIOPAC and RSPEC-R, PPGED-R systems respectively operating at 50Hz. Moving forward we want to compare the sensitivity of different EDA based sensors but it is outside the scope of this paper.^1^ Thermal imaging data were acquired through the FLIR Boson Thermal Camera and Seek Mosaic core cameras in tandem. The ASUS ROG Eye camera was used to record the RGB (Red, Green, Blue) video. This video data will be instrumental in developing and testing contactless monitoring systems such as remote Photoplethysmography (rPPG). However, the analysis of rPPG falls outside the scope of this paper. The software of the Multi-Spectral Reality Detector credibility assessment system developed by AI Seer Pte. Ltd. which also integrated the hardware apart from the BIOPAC, administered the test, synchronizing time-stamped data streams mentioned above and saving aligned data for subsequent analysis and predictions.

### 2.5 Participants

For our study, we targeted a sample size of 80 participants. To account for potential no-shows, equipment failures, and noisy data, we initially recruited 130 individuals. Ultimately, 118 experiments were successfully conducted, as 12 participants did not show up. Although the target of 80 participants was met, the rest of the sessions proceeded since the participants had already been scheduled. Recruitment was managed by AI Seer Pte. Ltd., leveraging Telegram posts to attract participants with reimbursement for time and travel of 30 SGD. Interested individuals responded via a Google Form, accessible through a QR code on the recruitment poster. Selected participants were then contacted via email to schedule their experiment slots. Prior to the experiment, a confirmation text message was sent to ensure attendance. Upon arrival, participants signed a Singapore Personal Data Protection Act (PDPA) form and a consent form (Appendix A). The demographics of our participants reflect the diverse Singaporean diaspora, in contrast to previous studies that often overlooked American minorities. The demographic breakdown of our participants by ethnicity and gender is detailed below in Table 3.

**Table 3.**
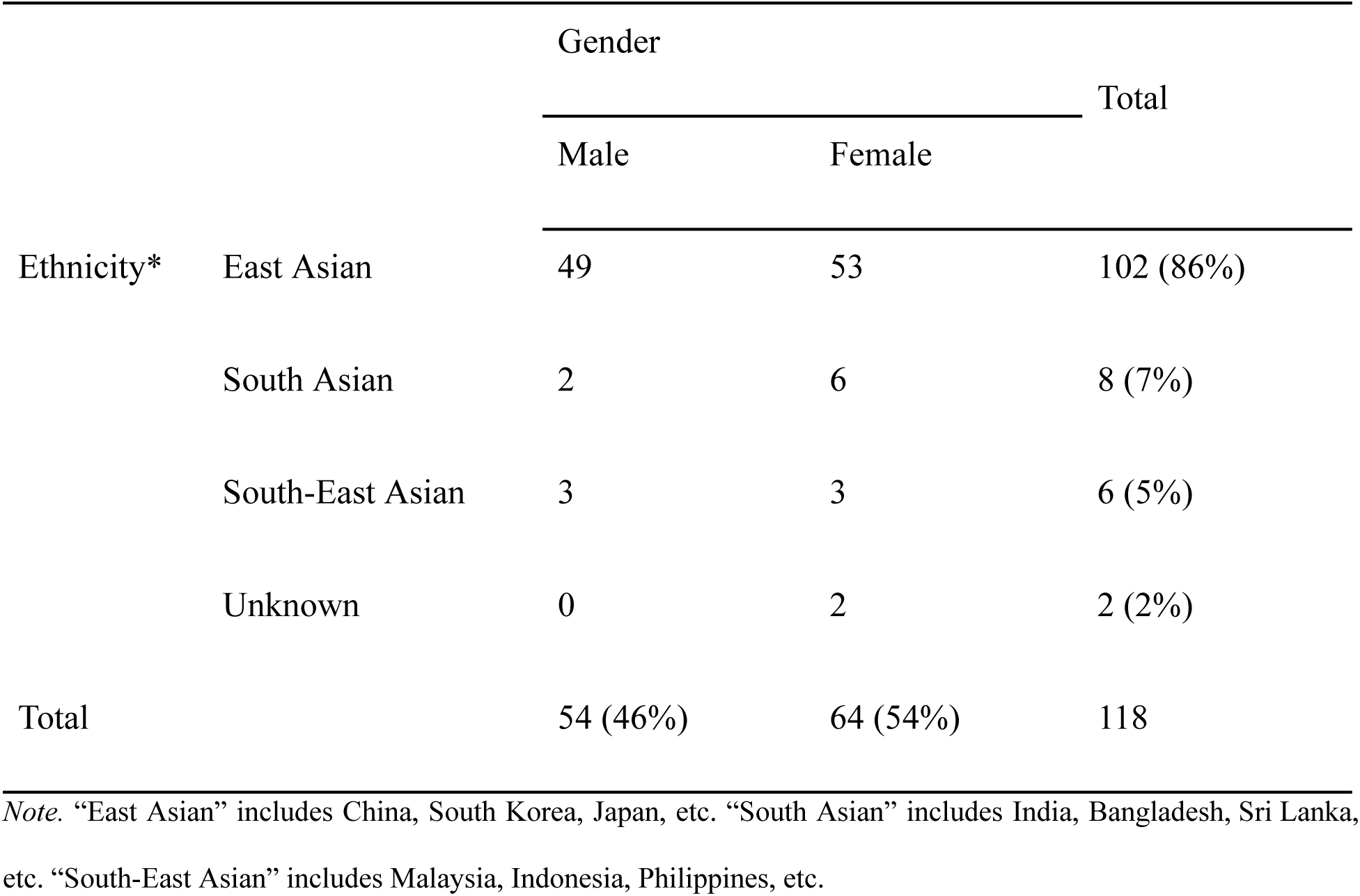
Demographics of Participants.

The study was conducted with 118 participants as shown in Table 3. Due to equipment failures, data was not collected from 12 participants but the participants were reimbursed for showing up for the experiment. Additionally, we excluded 15 datasets due to significant data corruption caused by noise, impacting the integrity of key physiological measures—pupil dilation, heart rate, thermal imaging, and EDA. The criteria for exclusion was that a dataset was deemed incomplete if more than 20% of data in any single channel for the required windows was obscured or distorted beyond our noise reduction capabilities. This threshold was determined through pilot testing, which showed that exceeding this level of noise substantially compromised the accuracy of our physiological analyses. Another exclusion criterion was answering more than 70% of the Moral Directed Lie questions wrongly. However, none of the participants did that and thus this was not a reason to exclude any of the collected data.

The sources of noise in our datasets were varied and stemmed from multiple factors, including equipment malfunctions, incorrect sensor placements, and participant movements. For instance, behaviors like participants looking away from the screen interrupted the collection of pupil and gaze data, while excessive finger movements disrupted EDA readings. Additionally, improper electrode placements skewed the electrocardiogram (ECG) data. These disturbances are typical in physiological studies and can significantly compromise the integrity of the data, particularly for precise metrics like pupil dilation monitoring. Consequently, examiners were asked to minimize their involvement in correcting these issues after the session had started to maintain the validity of the experimental conditions.

To preserve the scientific integrity of our analysis—especially given our interest in the interaction effects among these measures—it was crucial to ensure the completeness and consistency of the data across all channels for the entire dataset. Consequently, we opted to exclude these compromised datasets rather than attempt to impute or reconstruct the missing data. This decision was driven by the need for precision and accuracy in interpreting real-time physiological signals, which are vital for valid assessments of the differences between the Test and Control groups. After addressing these issues, we successfully gathered 91 complete datasets: 45 in the Test group and 46 in the Control group. Completeness was assessed based on whether all data channels provided acceptable, noise-free data.

### 2.6 Data Processing of dependent Variables

#### 2.6.1 The Present Research

This study addresses fundamental issues in deception detection by evaluating the effectiveness of physiological measures in distinguishing between deceptive and truthful individuals. Specifically, the research aims to determine whether physiological responses—such as pupil dilation, EDA, HR, and nose temperature—can reliably differentiate guilty participants from innocent ones. Based on prior research and theoretical frameworks, we proposed specific hypotheses (see preregistration <u>#151764 | AsPredicted</u>). In the preregistration, the term “critical question” was used for relevant questions, and “directed lie (DL)” was used for Moral Directed Lie (MDL) questions.

H1: Guilty participants will exhibit increased pupil dilation in their (relevant response - MDL response).

H2: Guilty participants will exhibit a greater EDA response in their (relevant response - MDL response).

H3: There will be greater HR deceleration for critical questions for guilty participants compared to innocent participants.

H4: There will be a greater decrease in nose temperature for guilty participants compared to innocent participants over the entire questioning period.

During the literature review conducted after the pre-registration, two additional hypothesis was formulated

H5: The difference in the decrease in heart rate from its initial peak to its minimum point between relevant questions and Moral Directed Lie questions will be greater for guilty participants compared to non-guilty participants.

H6: The difference in the RLL between relevant questions and Moral Directed Lie questions will be greater for guilty participants compared to non-guilty participants.

While H5 and H6 was not pre-registered, it was developed based on new insights gained during the literature review and will be explored in this study as an additional, non-pre-registered exploratory hypothesis.

experimental design with two conditions—guilty (intervention) and innocent (control)—was employed. Innocent participants were tasked with packing their luggage for a holiday abroad, including standard items such as a book, towel, and clothes. Guilty participants were required to conceal an additional contraband item and attempt to “smuggle” it across a simulated border. Following this, all participants were subjected to a moral directed-lie test and an Nback task. The Nback task will not be part of this paper. The key dependent variables are discussed in greater detail in the following section.

#### 2.6.2 H1: Pupil Area Under Curve Response

Pupil data was resampled at 30Hz for the entire session, with a focus on the area under the pupil response curve during the initial 8 seconds following question onset. To derive this, the pupil reading at question onset was subtracted from the subsequent response. The area under the curve was then computed for the 2 to 8-second timeframe. The initial moments after question onset often include artifacts or noise related to the transition period when the subject is first processing the question. Excluding this period reduces the impact of such artifacts, leading to more reliable and valid measurements of pupil response.

This approach also normalizes the responses by focusing on dynamic changes rather than absolute pupil size, making it easier to compare changes specifically induced by the question. Additionally, it ensures consistency with prior research practices and preregistration plans, strengthening the validity of the findings. This process was applied to both relevant and MDL questions, serving as the basis for comparison. The resulting values for all relevant responses and all moral lie responses were averaged separately. The feature under investigation was obtained by subtracting the average relevant response from the average moral lie response.

#### 2.6.3 H2: EDA Area Under Curve Response

EDA was resampled at a rate of 30Hz. The focus was on the area under the EDA curve during the initial 8 seconds following question onset. The EDA response during this period was extracted by subtracting the reading at the question onset from the subsequent values. The analysis extended from 0 to 8 seconds post-stimulus. The exclusion of the first 2 seconds in pupil data was specifically aimed at avoiding artifacts and noise associated with the initial cognitive processing phase. These artifacts may not be present in EDA data, so there was no compelling reason to exclude the initial 2 seconds for EDA analysis. The resulting EDA values for both relevant and MDL questions were separately averaged. Subsequently, the average EDA response to relevant questions was subtracted from the average response to moral lie questions, forming the EDA Response feature for examination.

#### 2.6.4 H3: Heart Rate Deceleration for Relevant Questions

The analysis focuses on heart rate deceleration between the fourth and eighth seconds following the presentation of relevant questions. This targeted approach highlights the distinct cardiac response associated with these questions, distinguishing it from broader analyses that examine heart rate dynamics across various question types.

In our previous in-house experiments, we found a significant difference in the heart rate deceleration slope when participants lied in response to relevant questions (test group) compared to when they answered truthfully (control group). Specifically, the deceleration slope was steeper in the test group, indicating a more pronounced cardiac response to deception. This analysis departs from the traditional CQT by focusing solely on the physiological response to relevant questions, without comparing it to other question types.

#### 2.6.5 H4: Changes in Facial Temperature

In our exploration of physiological responses, we conduct a detailed analysis of facial temperature changes across various regions throughout the experimental session. Leveraging an advanced 438-point 3D thermal facial mesh, we capture high-resolution thermal images of the face at both the beginning and the end of the session. This mesh enables precise temperature mapping across the entire face, facilitating a comprehensive examination of thermal variations.

To ensure accuracy despite subject movement, we flatten and project the thermal images onto a stable plane, allowing for consistent tracking of temperature changes over time. This process is crucial for maintaining the alignment of facial features and ensuring reliable comparisons of temperature data across the session. We focus particularly on the nose region, isolating and analyzing it by measuring the mean pixel intensity of the thermal image pixels for each frame. Unlike traditional approaches like the CQT, which typically compares physiological responses to specific stimuli, facial temperature changes do not occur rapidly within a short 15-second window.

Therefore, instead of examining temperature fluctuations in response to individual questions, our analysis measures the overall temperature change in each facial region from the start to the end of the session. The start of the session is defined as the first question, and the end of the session as the last question. The measurement is obtained by subtracting the temperature recorded at the start from that at the end.

By tracking these temperature deltas throughout the entire experimental period, we gain insights into how facial temperature evolves in response to the cumulative cognitive and emotional demands of the test. This methodology, inspired by our in-house experiments, underscores the importance of monitoring temperature changes over the full duration of a session rather than in reaction to isolated stimuli.

#### 2.6.6 H5: Heart Rate Peak-Decrease Response

To analyze heart rate variability (HRV), the neurokit2 library (https://pypi.org/project/neurokit2) was used for accurate ECG peak detection. The HRV was calculated by measuring the time interval in seconds between each ECG peak. To convert HRV data into heart rate (in beats per minute, BPM), the formula

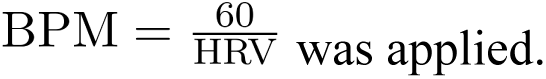

For heart rate response analysis, the beats per minute calculated data was resampled at a rate of 30Hz and normalized using z-score normalization across the entire session. The focus was particularly on the initial 15 seconds after the onset of a question. For this feature, the emphasis was on extracting distinctive heart rate dynamics during the initial 15 seconds following question onset. The maximum heart rate was determined within the first 5 seconds, representing the initial acceleration phase, while the minimum heart rate was identified between 5 and 15 seconds, reflecting the subsequent deceleration phase. The feature was computed by subtracting the minimum heart rate from the maximum heart rate, capturing the decrease in heart rate from the initial peak. The heart rate peak-decrease response to relevant questions was then subtracted from the peak-decrease response to moral directed lie questions, constituting the heart rate peak-decrease response. The selection of specific time windows (initial 5 seconds and subsequent 10 seconds) is motivated by empirical research on physiological responses to deception.

Studies, such as those by Verschuere et al. (2004) and Klein Selle et al. (2015), have shown that significant heart rate changes occur within seconds after a stimulus related to deception. The initial 5-second window captures the rapid heart rate increase due to sympathetic activation, a common response to stress and arousal associated with deception. The following 10-second window captures the deceleration phase, influenced by parasympathetic activation as the body attempts to restore homeostasis. This extended window captures the immediate orienting responses while also accommodating the prolonged deceleration associated with arousal inhibition. By comparing heart rates over this period, researchers can discern the contributions of OR and AI. The 15-second timeframe thus provides a balanced and inclusive perspective, enabling the identification of both transient and sustained physiological changes that are crucial for accurate detection of concealed information.

#### 2.6.6 H6: Respiratory Line Length Response

#### **a)** Original RLL

To quantify the changes in participants’ breathing patterns, RLL was calculated for each subject across defined time intervals. RLL measures the cumulative movement of the respiration waveform over time and was calculated as follows:

1. The respiration waveform was divided into small, consistent time segments (e.g., 0.1 seconds) over a 15-second window.
2. For each segment, the absolute difference in amplitude between consecutive data points was computed, providing the “moving distance.”
3. The total movement across all time segments in the window was summed.
4. The summed movement was divided by the number of segments within the time window to yield the average RLL, representing the total movement of the waveform during the window.

#### **b)** Weighted RLL

To account for variability in the lengths of respiratory cycles, a weighted RLL was also calculated as described by Matsuda (2011). This method mitigates potential bias by incorporating the duration of each respiratory cycle into the final RLL value:

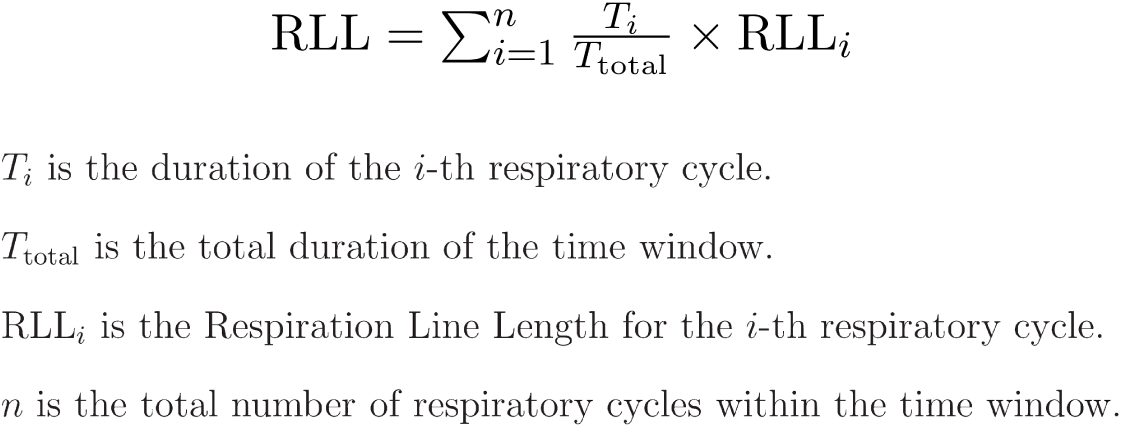

1. The respiration waveform was segmented into individual cycles by detecting peaks and troughs. Each respiratory cycle was defined as the interval between consecutive troughs.
2. The RLL was computed for each respiratory cycle by following the same steps used in the original RLL calculation, applied only to the data within each cycle.
3. The proportion of the total 15-second time window occupied by each respiratory cycle was calculated.
4. Each cycle’s RLL was multiplied by its corresponding time proportion. The sum of these weighted values yielded the final weighted RLL for the given time window.

This approach, as defined by Matsuda (2011), allowed us to address the variation in respiratory cycle lengths and provided a more representative analysis of the participants’ respiratory responses.

Although beyond the scope of this study, it is worth noting that respiration data could be used to probabilistically determine if an examinee is employing countermeasures, a technique often used by polygraph systems and examiners. For instance, advanced algorithms such as the Lafayette OSS-3 are designed to enhance polygraph accuracy by detecting such patterns (Liedetectortest.com, 2024). While this was not the focus of our current work, future studies might benefit from exploring the integration of such algorithms to look for such patterns before doing the analysis of data.

## 3. Results

### 3.1 Pupil Response

Figure 2 shows the results of pupil responses. From our analysis, distinct patterns emerged between guilty and non-guilty participants. For the group of 45 guilty participants, the mean pupil response revealed a significantly higher average pupil response when answering relevant questions for which the guilty participants were deceptive, compared to Moral Directed Lie Test (MDL) questions. In contrast, among the 46 non-guilty control participants, their pupil responses were lower during relevant questions for which they were being truthful and higher during MDL questions, where they were also being deceptive. This pattern aligns with our expectations, suggesting that lying in response to relevant questions imposes a greater cognitive load on guilty individuals. These findings underscore the differential impact of deception based on guilt status in cognitive processing.

**Figure 2.**
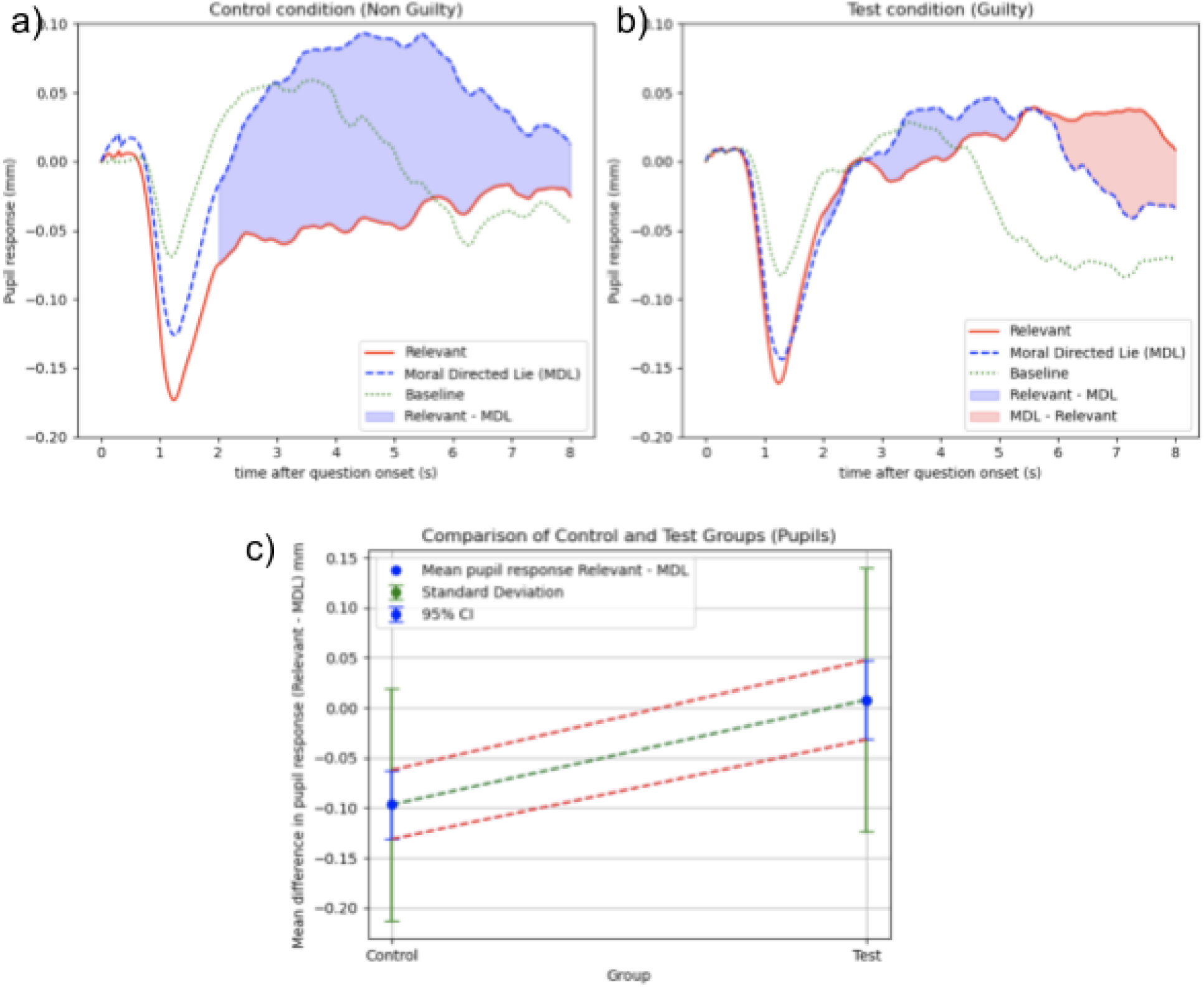
Pupil Response Results

Analyzing the difference in pupil response between MDL questions and relevant questions for both control (non-guilty) and test (guilty) groups reveals insightful patterns. For the control group, the mean difference in pupil response (relevant response - MDL response) was-0.09689, with a 95% confidence interval (CI) ranging from-0.1313 to-0.06247. This interval suggests that we can be 95% confident that the true mean difference in pupil response for the entire control population falls within this range. The negative values indicate that, on average, the pupil response during relevant questions is lower than during MDL questions, reflecting the increased cognitive load of lying in the MDL condition.

In contrast, the test group showed a mean difference in pupil response of 0.007677, with a 95% CI ranging from-0.03191 to 0.04276. This interval suggests that we can be 95% confident that the true mean difference in pupil response for the entire test population falls within this range. The values close to zero indicate that there is no significant difference in the pupil response between relevant and MDL questions for guilty participants. Between the control and test groups, the difference in pupil response (relevant response - MDL response) has a Cohen’s *d* of 0.85, indicating a large effect size. This large effect size underscores the substantial difference in cognitive load experienced by guilty and non-guilty participants when responding to relevant and MDL questions.

### 3.2 EDA Response

EDA responses are illustrated in Figure 3. They are distinguished between guilty and non-guilty participants, underscoring the differential impact of deception based on guilt status in cognitive processing. Specifically, there was a significant interaction between question type (relevant, directed moral lie) and group (control, test). This effect was driven by responses to relevant questions, which were significantly lower than MDL questions in the non-guilty group (mean difference in EDA response =-0.010602, 95% CI [-0.020065,-0.001139]), and significantly higher than MDL questions in the guilty group (mean difference in EDA response = 0.012924, 95% CI [0.001256, 0.024592]).

**Figure 3.**
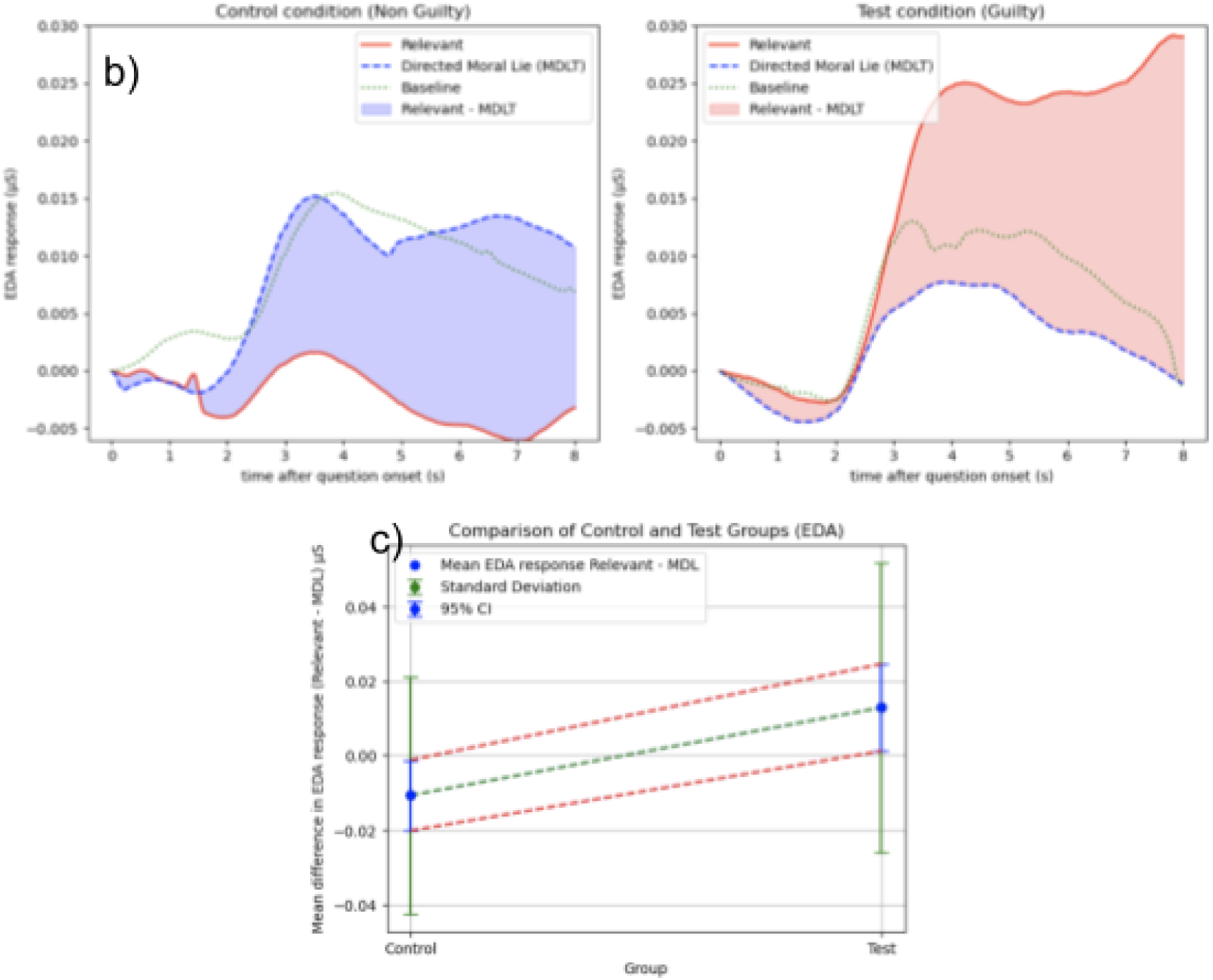
EDA Response Results

The negative values for the control group indicate that, on average, the EDA response during relevant questions is lower than during MDL questions, reflecting the increased cognitive load of lying in the MDL condition. In contrast, the positive values for the test group indicate that, on average, the EDA response during relevant questions is higher than during MDL questions, reflecting heightened arousal when answering deceitfully. Only the relevant question responses differed across groups, with a Cohen’s *d* of 0.66, indicating a medium to large effect size. This pattern aligns with our hypotheses and previous findings, highlighting the significant physiological response differences based on guilt status.

### 3.3 Heart Rate Response

Podlesny and Raskin (1978) found that HR responses to deception and truthfulness exhibited specific patterns. As shown in Figure 4, subjects showed an initial HR acceleration peaking at post-stimulus second 4, followed by a deceleration maximal around 11 seconds. Guilty subjects demonstrated greater HR deceleration in response to relevant questions compared to control questions, especially significant at 11 seconds. In line with the observations made by Podlesny and Raskin (1978), our experiment yielded results consistent with their findings.

**Figure 4.**
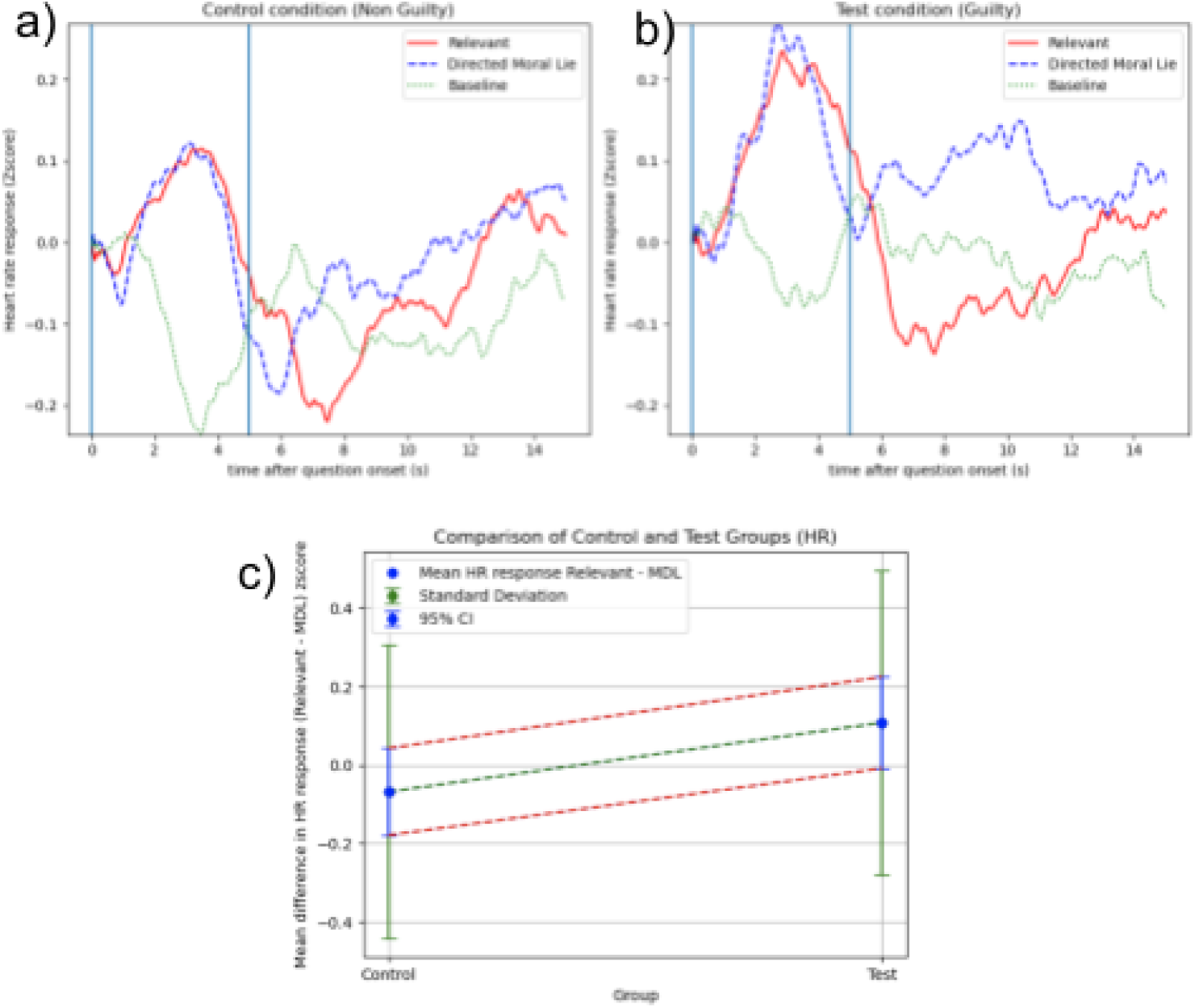
Heart Rate Response Results

In our study, both guilty and non-guilty participants exhibited a non-differential initial acceleration in heart rate, regardless of the type of question posed, peaking at around 4 seconds, whether it was a relevant question or an MDL question which for us worked as control questions. This initial increase in heart rate did not exhibit any noticeable disparity between the two participant groups, replicating the findings of Podlesny and Raskin.

Among the guilty participants, there was a notable reduction in heart rate when responding to relevant questions compared to comparison MDL questions. This outcome suggests that guilty individuals experienced a more pronounced relaxation or decrease in heart rate upon concluding their deceptive responses to relevant questions. In contrast, this distinctive pattern was not observed in the non-guilty group, consistent with the findings of the original study by Podlesny and Raskin (1978). Overall, the heart rate patterns and the interaction between guilt and question type were similar in both studies, with only minor differences.

A difference between our study and the findings of Podlesny and Raskin (1978) lies in the timing of the lowest heart rate decrease post-stimulus. They suggested that the lowest heart rate decrease occurred at 11 seconds post-stimulus. However, our study revealed a different pattern. We observed that the lowest heart rate occurred within a range of 5 to 15 seconds after an initial acceleration phase, which itself occurred within the first 5 seconds post-stimulus and not strictly at the fourth second. Instead of selecting specific points, like the 4th and 11th seconds which seem to be motivated by the mean plots in the Podlesny & Raskin (1978) study for maximum and minimum heart rates, we defined a time range of 0 to 5 seconds for maximum acceleration and 5 to 15 seconds for minimum deceleration. This approach yielded better results, as the peak acceleration and subsequent deceleration times vary between participants. By calculating these values within a window, we ensure a more accurate representation of the full extent of heart rate acceleration and subsequent deceleration where the delta between this is what differentiates for guilt.

Furthermore, our experiment included an additional hypothesis regarding the rate of change in heart rate during the deceleration phase (4-8 seconds). In a previous in-house experiment, we found that the slope of heart rate deceleration for relevant questions exhibited differential abilities, specifically distinguishing between guilty and non-guilty participants. However, our current findings did not support this hypothesis. We did not observe any significant differences in the rate of heart rate change between the two groups during this critical period.

### 3.4 Respiration Response

The Original RLL (Relevant - MDL) for the Test group had a mean of-1.14 with a 95% confidence interval (CI) of [-2.41, 0.13], while the Control group had a mean of 0.13 with a 95% CI of [-0.95, 1.21]. With a p-value of 0.14, this result indicates no significant difference between the groups. Similarly, the Weighted RLL (Relevant - MDL) for the Test group had a mean of-0.16 with a 95% CI of [-0.39, 0.07], and the Control group had a mean of 0.05 with a 95% CI of [-0.22, 0.33]. The p-value here was 0.24, again showing no statistically significant difference between the groups. Overall, the RLL (Relevant - MDL) measures—both Original and Weighted—did not reveal any significant differences between the Test and Control groups. However, it’s worth noting that the Test group had consistently negative RLL values for both measures, suggesting the expected directionality: guilty participants had shorter RLLs for relevant questions compared to MDL questions. While this pattern aligns with expectations, it was not statistically significant.

RLL is often considered less reliable than other physiological signals, such as EDA, in deception detection studies (Synnott, Dietzel & Ioannou, 2015). The evidence suggests that EDA offers more consistent and accurate insights into physiological responses compared to RLL. This is in line with the findings of this paper.

### Experimental Respiration Calculations

We experimented with a new method to better understand RLL responses by tracking respiratory amplitude over a 15-second window, which offered a more precise identification of shallow breathing points. This feature was not pre-registered but we believe can be valuable to explore further to compare it to the traditional methods of calculating RLL. Traditional RLL methods didn’t replicate in our study, so we employed the neurokit2 library to process the raw respiration signal, accurately detecting respiration cycles.

We handled the respiration depth of each cycle independently along the time axis. We measured respiration depth and amplitude by calculating the vertical distance between the peak and trough of each cycle, representing the depth of each breath. For each respiration cycle, the measured depth was kept constant throughout the duration of that cycle. This approach allowed for a more accurate representation of the respiration depth across different time periods. By standardizing this data with z-score normalization across the entire session, we captured detailed changes in respiration depth following question onset. The respiratory amplitude response was then derived by subtracting the baseline depth at question onset from 15 seconds of depth measure post question onset. Comparing these changes between relevant and morally directed lie questions revealed distinct respiratory patterns.

In Figure 5, we present the variations in respiratory amplitude from the onset of questioning. The data reveals a consistent pattern of respiratory suppression following the initiation of questions across all question types. Notably, there is a minimal divergence in the respiratory amplitude trends for non-guilty participants between the relevant and MDLT questions. This indicates a relatively uniform respiratory response regardless of the question type. Conversely, the data for guilty participants exhibits a more pronounced difference. Here, we observe a significant decrease in respiratory amplitude, particularly in response to relevant questions in comparison to MDL. The mean difference in respiratory depth between relevant and MDL questions yielded a p-value of 0.045 and a Cohen’s d of 0.43, indicating a medium effect size. This feature emerged during data analysis and we are not aware of it being a common deception cue used by polygraphers or that it is currently in the literature, suggesting that further research is necessary to explore the replicability and validity of this measure.

**Figure 5.**
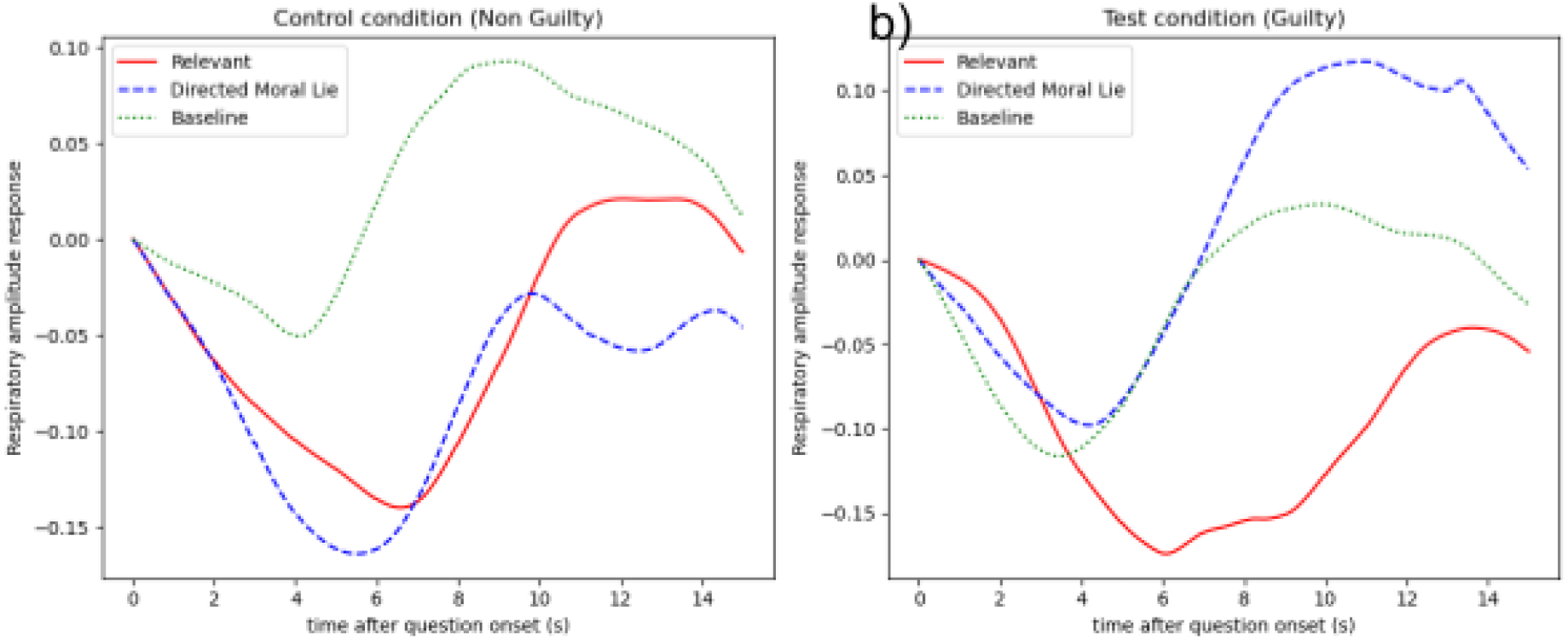
Experimental Feature of Respiration Depth Amplitude

## 4. Discussion

An analysis of our results reveals that pupil response had the most substantial impact in differentiating between truthful and deceptive conditions, as evidenced by its large Cohen’s *d* and high statistical significance highlighted in Table 4. Heart rate response also shows a meaningful but smaller effect, indicating it still contributes to distinguishing between the groups. EDA response similarly demonstrates a significant and moderately strong effect, further validating its role in the differentiation. However, the change in nose temperature, heart rate deceleration as well as RLL was found to be non-significant in this dataset.

**Table 4.** Outcomes of the Tested Features.

| Feature | $p$ | Cohen's $d$ | $t$ |
| --- | --- | --- | --- |
| Pupil Response (H1) | 0.0001 | 0.853 | 4.016 |
| EDA Response (H2) | 0.0023 | 0.668 | 3.144 |
| Heart Rate Response (H5) | 0.0297 | 0.469 | 2.210 |
| Heart rate deceleration (H3) | 0.451 | 0.160 | 0.764 |
| RLL original (H6 a) | 0.141 | 0.312 | 1.486 |
| RLL weighted (H6 b) | 0.242 | 0.247 | 1.179 |
| Respiratory depth (H6 experimental) | 0.045 | 0.427 | 2.037 |
| Nose Temperature (H4) | 0.4390 | 0.165 | 0.777 |

Additionally, regression analyses were conducted using different combinations of the significant responses— pupil response (H1), heart rate response (H5), and EDA response—(H2) to determine which combination best explains the variance in differentiating between truthful and deceptive conditions. The results can be seen in Table 5 below. This helped to identify the most predictive variables and refine our understanding of their individual and combined contributions to the model.

**Table 5.**
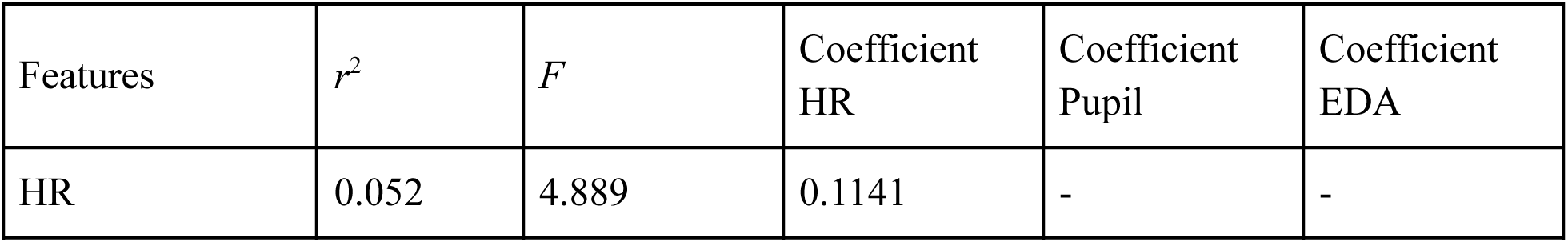
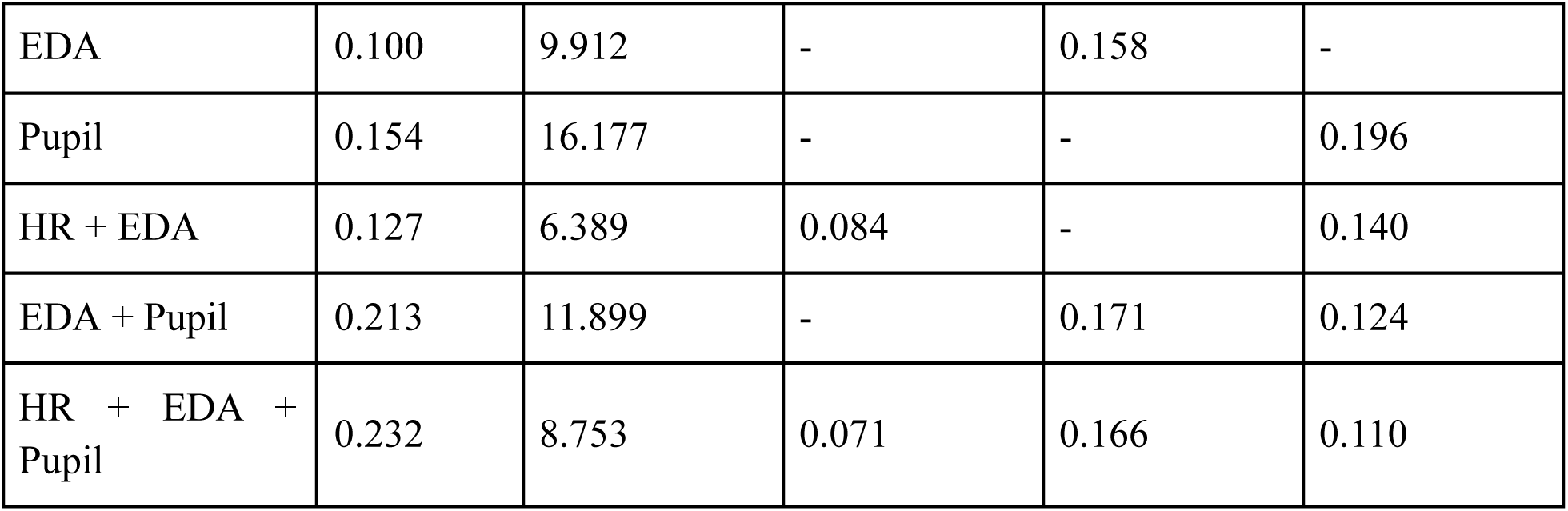
Regression Model Statistics of Various Feature Combinations.

The analysis of various regression models using HR, EDA, and pupil response as predictors for differentiating between truthful and deceptive conditions reveals nuanced insights into their individual and combined contributions. The model with only HR explains 5.2% of the variance (*R*-squared = 0.052), with a modest contribution from HR (coefficient = 0.1141) and an *F*-statistic of 4.889, indicating a significant but weak predictive power. In contrast, the EDA-only model improves the variance explained to 10.0% (*R*-squared = 0.100) with a stronger effect (coefficient = 0.158) and a higher *F*-statistic of 9.912, suggesting that EDA is a more robust predictor than HR alone.

Pupil response, however, emerges as the most powerful single predictor, explaining 15.4% of the variance (*R*-squared = 0.154) with a coefficient of 0.196 and an *F*-statistic of 16.177, highlighting its significant role in distinguishing between the conditions. When combining HR and EDA, the model’s explanatory power increases slightly to 12.7% (*R*-squared = 0.127), with coefficients of 0.084 for HR and 0.140 for EDA, but the F-statistic drops to 6.389, indicating a reduced relative significance of the predictors.

The model combining EDA and pupil explains a substantial 21.3% of the variance (*R*-squared = 0.213), with both predictors contributing significantly (coefficients of 0.171 and 0.124, respectively) and an *F*-statistic of 11.899, showing a robust combined effect. The full model, incorporating HR, EDA, and pupil response, achieves the highest R-squared value of 0.232, explaining 23.2% of the variance, with coefficients of 0.071 for HR, 0.166 for pupil, and 0.110 for EDA. However, the *F*-statistic of 8.753, while still significant, is lower than that of the pupil-only model, suggesting that the added complexity of including HR and EDA does not proportionately enhance the model’s overall significance. These findings suggest that while the combination of all three predictors provides the most comprehensive model in terms of variance explained, pupil response alone or in combination with EDA offers a more balanced approach, maximizing predictive power while maintaining model simplicity and statistical significance.

Moving forward, we explored whether significant interactions exist between HR, EDA, and pupil response in predicting whether a participant is truthful or deceptive. To achieve this, interaction terms were created by multiplying the values of the involved predictors (e.g., HR * EDA) to capture potential combined effects. These interaction terms were then included in regression models alongside the individual predictors.

For each model, we calculated the p-values associated with the interaction terms to determine their statistical significance. Specifically, the following steps were taken:

1. **Model Development:** Regression models were built with interaction terms such as HR * EDA, HR * Pupil, EDA * Pupil, and the three-way interaction HR * EDA * Pupil
2. **Statistical Testing:** Each model was fitted using Ordinary Least Squares (OLS) regression, and the p-values for the interaction terms were extracted
3. **Significance Assessment:** The p-values were compared to a significance level to assess whether the interaction terms significantly contributed to explaining the variance in the dependent variable (truthful vs. deceptive condition)

The analysis shown in Table 6 reveals that none of the interaction terms between HR, EDA, and pupil response are statistically significant, with all p-values exceeding the typical threshold of 0.05. This indicates that the effects of HR, EDA, and pupil response on predicting deception do not significantly depend on the levels of the other measures. Therefore, the individual effects of these physiological measures are more relevant and do not benefit from including interaction terms in this context. The findings suggest that the model’s explanatory power is primarily driven by the individual contributions of HR, EDA, and pupil response, rather than by their combined interactions. AI Seer is also working on a paper that discusses its classification model and its sensitivity settings that can allow the system to be used in various contexts.^2^

**Table 6.**
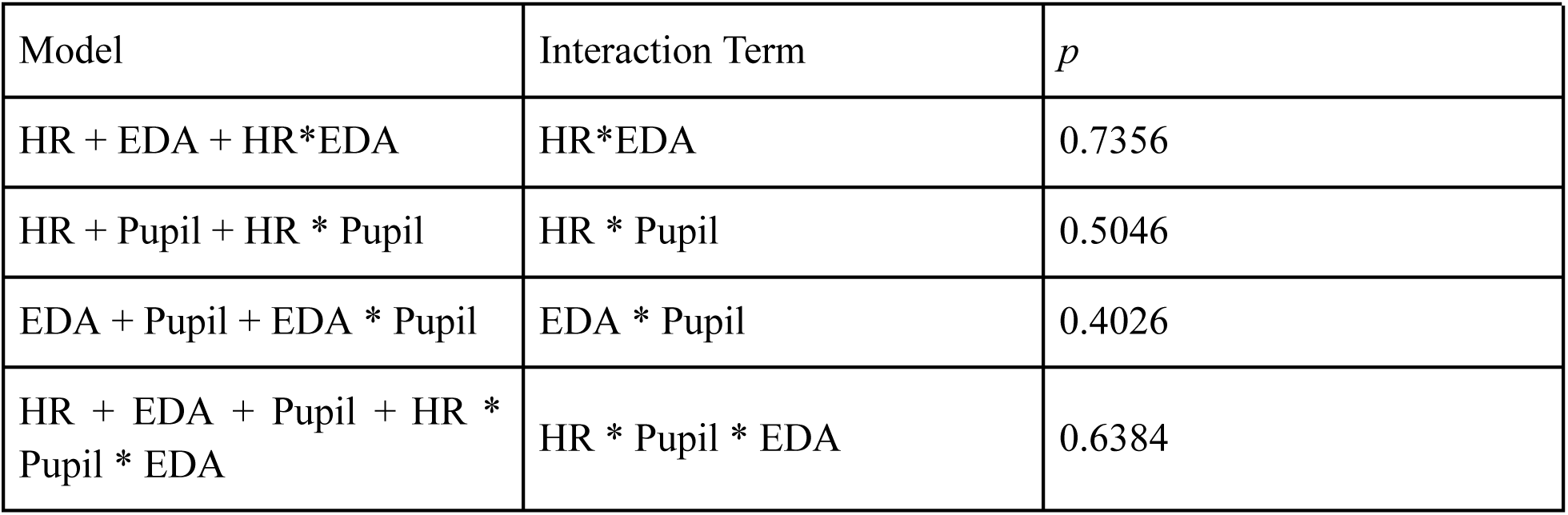
Interaction Effect of Different Responses.

| Model | Interaction Term | *p* |
| --- | --- | --- |
| HR + EDA + HR*EDA | HR*EDA | 0.7356 |
| HR + Pupil + HR * Pupil | HR * Pupil | 0.5046 |
| EDA + Pupil + EDA * Pupil | EDA * Pupil | 0.4026 |
| HR + EDA + Pupil + HR * Pupil * EDA | HR * Pupil * EDA | 0.6384 |

**Table 7.**
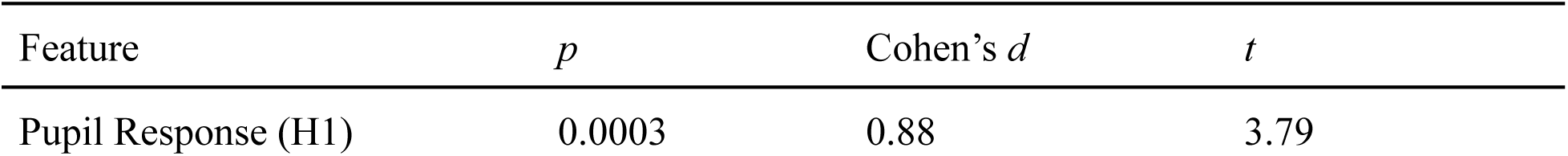
Regression Model Statistics of Pupil Response using traditional Directed Lie Paradigm (n=74, 37 test, 37 control)

## 5. Conclusion

The findings reveal that pupil response emerges as the most robust and significant predictor of deception within the framework of the CQT. Incorporating EDA as a secondary predictor further enhances the model’s explanatory power. Although HR remains relevant, its contribution is comparatively modest, offering limited additional value when combined with pupil response and EDA. Meanwhile, changes in nose temperature were found to be statistically non-significant in this dataset, indicating that this measure may not be a reliable predictor of deception in the contexts examined The absence of significant interaction effects between these physiological measures suggests that their individual contributions are currently more impactful when considered separately. However, this highlights the need for further research, potentially exploring alternative questioning techniques, to better understand how these physiological indicators might interact under varying conditions and enhance deception detection in CQT scenarios.

In our previous study using the traditional Directed Lie paradigm (n = 74; Cohen’s d = 0.88, t = 3.79), we replicated the expected pupil-dilation effect but received frequent reports of confusion about the question format. In the present study, the Moral Directed Lie Test (MDLT) produced a highly similar effect size (d = 0.85, t = 4.02 with n = 91), indicating that the new moral-control paradigm preserves sensitivity to deception-related pupil responses while offering clearer, more universally interpretable instructions. Although the slightly larger *t*-statistic in the MDLT study is largely attributable to the increased sample size rather than a stronger underlying effect, the comparable effect sizes suggest that the MDLT can serve as a viable alternative to the traditional DLT.

### 5.1 Limitations of Current Research and Future Directions

While the current study provides valuable insights into the physiological indicators of deception, several limitations must be acknowledged. First, the scope of this research was confined to a specific set of physiological measures— pupil response, EDA, facial temperature change and HR— within the context of the CQT. While these measures have shown promise, the exclusion of other potential indicators, such as blood volume pressure, limits the comprehensiveness of our findings. Future research should consider integrating a broader range of physiological and behavioral measures to create a more holistic model of deception detection.

Another limitation pertains to the generalizability of the findings. The study was conducted in a controlled experimental setting, which offers a high degree of control and consistency, however, it may not fully capture the complexities of real-world deception. Participants might behave differently in more naturalistic environments, where stakes are higher, or when deception involves more nuanced social interactions. Future research should aim to validate these findings in more ecologically valid settings such as in real-world law enforcement, security screenings or corporate settings, to ensure that the results are applicable beyond the laboratory.

Additionally, the lack of significant interaction effects between the physiological measures suggests that our current models may not fully capture the complexity of how these indicators interact in predicting deception. Future studies should explore alternative statistical approaches or machine learning techniques to identify potential nonlinear interactions or combined effects that traditional regression models might overlook. By leveraging these advanced techniques, future research could uncover hidden patterns or relationships between physiological indicators that could significantly enhance the accuracy of deception detection.

Moreover, while the study identified pupil response as the most significant predictor of deception, the current approach does not fully explore the temporal dynamics of physiological responses. Deception is not a static process, it unfolds over time, with different indicators becoming more or less salient to varying stages of the questioning process. Future research could employ time-series analysis or dynamic modeling techniques to better understand these temporal patterns, offering a more detailed and granular understanding of how physiological responses evolve during deceptive behavior.

Lastly, the current research relied on a single type of questioning technique (CQT). However, different questioning methods, such as the Concealed Information Test (CIT) or Guilty Knowledge Test (GKT), may elicit different physiological responses and could potentially interact with the measures investigated in this study. Exploring these alternative questioning techniques in future research could determine whether they can enhance the detection of deception when combined with the physiological indicators identified in this study.

In conclusion, while this research contributes to the understanding of physiological indicators of deception, it also opens up several avenues for future investigation. By addressing these limitations and expanding the scope of inquiry, future studies have the potential to build on the foundation laid by this research to develop more accurate and reliable deception detection methods. Such advancements could have profound implications in various fields including law enforcement and beyond, ultimately enhancing truth-seeking processes in diverse contexts.

## 6. Declaration Abbreviations

● AI: Artificial Intelligence
● ANS: Autonomic Nervous System
● CIT: Concealed Information Test
● CQT: Comparison Question Test
● ECG: Electrocardiography
● EDA: Electrodermal Activity
● GKT: Guilty Knowledge Test
● GSR: Galvanic Skin Response
● HR: Heart Rate
● HRV: Heart Rate Variability
● IR: Infrared
● MDL: Moral Directed Lie
● MDLT: Moral Directed Lie Test
● N-back: A continuous performance task used to assess working memory
● PD: Pupil Diameter
● PDPA: Personal Data Protection Act
● PIEC: Parkway Independent Ethics Committee
● PNS: Parasympathetic Nervous System
● PPG: Photoplethysmography
● RLL: Respiratory Line Length
● RSA: Respiratory Sinus Arrhythmia
● RSP: Respiration
● SNS: Sympathetic Nervous System

## Ethics Approval and Consent to Participate

The study was conducted in accordance with the ethical standards outlined in the Declaration of Helsinki and was approved by the Parkway Independent Ethics Committee (PIEC) under IRB reference number PIEC/2022/062. Written informed consent was obtained from all participants prior to their inclusion in the study. Participants were fully informed about the nature of the study, including the use of deception, and were assured of their right to withdraw at any time without penalty.

## Consent for publication

Not Applicable

## Availability of Data and Materials

The processed datasets generated is made available and analyzed during the current study are not publicly available due to confidentiality agreements but are available from the corresponding author on reasonable request. The processed data is available on https://doi.org/10.7910/DVN/TQUXVG.

## Authors’ Contributions

● **A M Shahruj Rashid**: Conceptualization, Methodology, Data Collection, Data Analysis, Writing—Original Draft, Correspondence.
● **Bryan Carmichael**: Literature Review, Data Collection, Writing—Review & Editing.
● **Charlize Su**: Data Collection, Writing—Review & Editing.
● **Keming Shi**: Data Collection, Writing—Review & Editing.
● **Keefe Lim**: Data Collection, Writing—Review & Editing.
● **Senthil Kumar Poorvika**: Data Collection, Literature Review, Writing—Review & Editing.
● **Ngok Jeun Wan**: Data Collection.
● **Eshaan Govil**: Writing—Review & Editing.
● **Dennis Yap**: Supervision, Project Administration, Funding Acquisition, Writing—Review & Editing, Correspondence.

All authors read and approved the final manuscript.

## Supporting information

Appendix A PDPA and Consent Form

## Acknowledgements

We would like to express our gratitude to the polygraph experts, Defense Tech R&D officials and academics who generously provided us with their valuable advice and insights, any errors or omissions remain our own. The authors declare no other conflicts of interest. We extend our gratitude to Parkway Health for providing Institutional Review Board (IRB) approval for this study, to the participants who took part in the research, and to AI Seer Pte. Ltd. for providing the necessary facilities and resources

## Footnotes

1 AI Seer Pte. Ltd.,Investigation of Correlation and Sensitivity: BIOPAC EDA vs. Arduino System EDA and BIOPAC Respiration vs Thermal Camera Based Respiration. Work in progress.

2 AI Seer Pte. Ltd., “Development and Sensitivity Analysis of the Facticity Classification Model: Adapting Sensitivity Settings for Diverse Use Cases,” work in progress.

