## Appendix A PDPA and Consent Form for "Evaluating Physiological Indicators in Detecting Deception and Truthfulness Using the Comparison Question Test"

AI Seer Pte. Ltd.
UEN: 201942044R
24 St. Patrick's Road
Singapore 424146

### DATA PROTECTION NOTICE

This Data Protection Notice ("Notice") sets out the basis which AI Seer Pte. Ltd. ("AI Seer", "we", "us", or "our") may collect, use, disclose or otherwise process personal data (including personal data of our customers, study subjects, and employees) in accordance with the Personal Data Protection Act 2012 (No. 26 of 2012) ("PDPA"). This Notice applies to personal data in our possession or under our control, including personal data in the possession of organisations which we have engaged (if any), or have engaged us, to collect, use, disclose or process such personal data for the purposes set out in this Notice.

#### PERSONAL DATA

1. As used in this Notice:

   "**biometrics**" include, but is not limited to, (a) heart rate; (b) respiration rate; (c) eye and pupil images and pupil dilations; (d) shoulder movements; (e) facial expressions and facial movements including blinking, micro-expressions and movements of the mouth, lips, and chin; (f) voice and speech patterns, pitch and tone; and (g) other available biometric information captured or collected by our good or service;

   **"customer"** means a corporate entity, organization or government agency who (a) has contacted us through any means to find out more about any goods or services we provide, (b) may, or has, entered into a contract with us for the supply of any goods or services by or to us; or (c) is a paid or unpaid user of our goods or services;

   **"personal data"** means data, whether true or not, about a customer's officer, director, employee, or representative, or a subject who can be identified: (a) from that data; or (b) from that data and other information to which we have or are likely to have access;

   "**subject**" means an individual who has participated in our research or usability study involving the collection, use and/or processing of user information and biometrics (which may include personal data).

2. Depending on the nature of your interaction with us, some examples of personal data which we may collect from you may include your name and identification information such as your NRIC number, contact information such as your address, email address or telephone number, nationality, gender, date of birth, photographs, biometrics and other audio-visual information.

3. Other terms used in this Notice shall have the meanings given to them in the PDPA (where the context so permits)

#### COLLECTION, USE AND DISCLOSURE OF PERSONAL DATA

4. Your personal data is collected when:

   (a) it is provided to us voluntarily by you directly or via a third party who has been duly authorised by you to disclose your personal data to us (your "authorised representative") after:

   (i) you (or your authorised representative) have been notified of the purposes for which the data is collected, and

   (ii) you (or your authorised representative) have provided written consent to the collection and usage of your personal data for those purposes, or

(b) collection and use of personal data without consent is permitted or required by the PDPA or other laws. We shall seek your consent before collecting any additional personal data and before using your personal data for a purpose which has not been notified to you (except where permitted or authorised by law).

5. We may collect and use your personal data for any or all of the following purposes:

(c) performing obligations in the course of or in connection with the study;

(d) verifying your identity;

(e) responding to, handling, and processing queries, requests, applications, complaints, and feedback from you;

(f) managing your relationship with us;

(g) processing payment or credit transactions;

(h) inviting you to participate in future studies

(i) complying with any applicable laws, regulations, codes of practice, guidelines, or rules, or to assist in law enforcement and investigations conducted by any governmental and/or regulatory authority;

(j) any other purposes for which you have provided the information;

(k) any other incidental business purposes related to or in connection with the above;

(l) improving our algorithms for commercial and non-commercial purposes;

(m) business improvements, in order to carry out operational efficiency and service improvements, or to develop, enhance, or evaluating the effectiveness of our goods, service, or equipment; and

(n) for research purposes.

6. We may disclose your personal data:

(a) to such customer(s) or parties that may be stated in the relevant study consent form. Such customer may have the right to ownership of the data captured during the study, including biometrics;

(b) where such disclosure is required for incidental business or research purposes, or for performing obligations in the course of or in connection with our provision of the goods or services requested by you; or

(c) for purposes of marketing and informing third parties about our product; or

(d) to third party service providers, agents and other organisations we have engaged to perform any of the functions listed in clause 5 above for u; or

(e) if permitted under applicable law. In the event that results of our research are published, we will publish the results in a form that does not identify you.

7. The purposes listed in the above clauses may continue to apply even in situations where you have no relationship with us, or your relationship with us (for example, pursuant to a contract) has been terminated or altered in any way, for a reasonable period thereafter.

#### WITHDRAWING YOUR CONSENT

8. The consent that you provide for the collection, use and disclosure of your personal data will remain valid until such time it is being withdrawn by you in writing. You may withdraw consent and request us to stop using and/or disclosing your personal data for any or all of the purposes listed above by submitting your request in writing or via email to our Data Protection Officer at the contact details provided below.

9. Upon receipt of your written request to withdraw your consent, we may require reasonable time (depending on the complexity of the request and its impact on our relationship with you) for your request to be processed and for us to notify you of the consequences of us acceding to the same, including any legal consequences which may affect your rights and liabilities to us. In general, we shall seek to process your request within ten (10) business days of receiving it.

10. Whilst we respect your decision to withdraw your consent, (a) if you are a customer, please note that depending on the nature and scope of your request, we may not be in a position to continue providing our goods or services to you, and (b) if you are a subject, to the extent that we still own and control your personal data, we reserve the right avail to exceptions to consent under applicable law as at the date of your written request or to pass on your request to the controller of your personal data, and we shall, in such circumstances, notify you before completing the processing of your request. Should you decide to cancel your withdrawal of consent, please inform us in writing in the manner described in clause 8 above.

11. Please note that withdrawing consent does not affect our right to continue to collect, use and disclose personal data where such collection, use and disclose without consent is permitted or required under applicable laws.

#### ACCESS TO AND CORRECTION OF PERSONAL DATA

12. If you are a customer or a subject and you wish to make (a) an access request for access to a copy of the personal data which we hold about you or information about the ways in which we use or disclose your personal data, or (b) a correction request to correct or update any of your personal data which we hold about you, you may submit your request in writing or via email to our Data Protection Officer at the contact details provided below.

13. Please note that a reasonable fee may be charged for an access request. If so, we will inform you of the fee before processing your request.

14. To the extent that we own and control your personal data, we will respond to your request as soon as reasonably possible. Should we not be able to respond to your request within thirty (30) days after receiving your request, we will inform you in writing within thirty (30) days of the time by which we will be able to respond to your request. If we are unable to provide you with any personal data or to make a correction requested by you, we shall generally inform you of the reasons why we are unable to do so (except where we are not required to do so under the PDPA).

#### PROTECTION OF PERSONAL DATA

15. To safeguard your personal data from unauthorised access, collection, use, disclosure, copying, modification, disposal or similar risks, we have introduced appropriate

administrative, physical and technical measures such as up-to-date antivirus protection, encryption and the use of privacy filters to secure all storage and transmission of personal data by us, and disclosing personal data both internally and to our authorised third party service providers and agents only on a need-to-know basis.

16. You should be aware, however, that no method of transmission over the Internet or method of electronic storage is completely secure. While security cannot be guaranteed, we strive to protect the security of your information and are constantly reviewing and enhancing our information security measures.

#### ACCURACY OF PERSONAL DATA

17. We generally rely on personal data provided by you (or your authorised representative). In order to ensure that your personal data is current, complete and accurate, please update us if there are changes to your personal data by informing our Data Protection Officer in writing or via email at the contact details provided below.

#### RETENTION OF PERSONAL DATA

18. We may retain your personal data for (a) the period stated in the study consent form; (b) as long as it is necessary to fulfil the purpose for which it was collected, or (c) as required or permitted by applicable laws, whichever is the longest.

19. We will cease to retain your personal data, or remove the means by which the data can be associated with you, as soon as it is reasonable to assume that such retention no longer serves the purpose for which the personal data was collected, and is no longer necessary for legal or business purposes.

#### TRANSFERS OF PERSONAL DATA OUTSIDE OF SINGAPORE

20. We generally do not transfer your personal data to countries outside of Singapore. However, if we (or our customer(s) that controls the personal data collected) do so, we or our relevant customer (depending on which party owns and controls the personal data collected) shall obtain your consent for the transfer to be made and to the extent that we own and control your personal data, we will take steps to ensure that your personal data continues to receive a standard of protection that is at least comparable to that provided under the PDPA.

#### DATA PROTECTION OFFICER

21. You may contact our Data Protection Officer if you have any enquiries or feedback on our personal data protection policies and procedures, or if you wish to make any request, in the following manner: Veronica Sng, 79 Ayer Rajah Crescent, #02-18, Singapore 139955,, +65 90608493 or Dennis Yap, 79 Ayer Rajah Crescent, #02-18, Singapore 139955,, +65 83050508

#### EFFECT OF NOTICE AND CHANGES TO NOTICE

22. This Notice applies in conjunction with any other notices, contractual clauses and consent clauses that apply in relation to the collection, use and disclosure of your personal data by us.

23. We may revise this Notice from time to time with or without any prior notice. You may determine if any such revision has taken place by referring to the date on which this Notice was last updated. Your continued use of our services or non-withdrawal of consent constitutes your acknowledgement and acceptance of such changes.

AI Seer Pte. Ltd.
UEN: 201942044R
24 St. Patrick's Road
Singapore 424146

Effective date : 07/02/2023
Last updated : 07/02/2023

ACKNOWLEDGEMENT AND CONSENT

I acknowledge that I have read and understood the above Data Protection Notice, and consent to the collection, use and disclosure of my personal data by AI Seer Pte. Ltd. for the purposes set out in the Notice.

Name (per NRIC): ______________________________________________

Signature: ____________________________________________________

Date: ________________________________________________________

AI Seer Pte. Ltd.
UEN: 201942044R
24 St. Patrick's Road
Singapore 424146

### CONSENT TO PARTICIPATE IN USABILITY STUDY
### Deception Detection with an Automated Interviewing System – Alibi Game

#### Introduction

You are invited to participate in a study conducted by AI Seer Pte. Ltd. This research study is being conducted in order to develop an Automated Deception Detection Interview System, including its analysis algorithm.

This sheet provides you with information about the research study. The researcher in charge of this study will describe this research to you and answer all of your questions. Please read the information below carefully and ask any questions about the study as you like before deciding whether or not to take part. The study staff can explain words or information that you do not understand. If you decide to take part in this study, you are required to sign your name at the end of this form and date it. A copy of this form will be provided to you upon request.

#### Purpose of the Study

The main objective of this study is for AI Seer Pte. Ltd. to (a) evaluate the effectiveness of the application and its underlying data analytics; and (b) validate the sensitivity of the sensors in collecting physiological data from participants, in developing a credibility assessment application.

#### What will happen during the study?

Your participation in this study will require 20 minutes to 40 minutes of your time. You will be required to participate in computerized tasks, during which you may be required to respond in a non-truthful manner.

#### Subject Participation

We estimate that around 90 participants will enroll in this study. Participants' age must range from 21 years old to 65 years old at the time of enrollment. Participants should have no history of cognitive impairment or any conditions that could significantly impact their ability to comprehend and engage with the study materials. Participants must also have no history of psychological disorders or psychiatric illnesses. Lastly, participants must possess a minimum level of proficiency in the English language.

#### Data Collected

As part of the study, AI Seer Pte. Ltd. will collect the following data:

- Demographic information (Name, Gender, Age, Race, and Language proficiency)
- Contact details: handphone number and email
- Physiological data (including but not limited to: (a) heart rate; (b) respiration rate; (c) eye and pupil images and pupil dilations; (d) shoulder movements; (e) facial expressions and facial movements including blinking, micro-expressions and movements of the mouth, lips, and chin; (f) voice, speech patterns, pitch, and tone), and (g) skin conductance
- Video and audio data of study participants

**Cost and Payments related to Participation in the Study**

You will receive $20 in remuneration for your participation. No expenses will be incurred by you as a result of your participation in this study.

**Potential Benefits**

Other than the incentive provided, this study is not expected to bring you direct benefits outside of this opportunity. Your participation, however, will be of considerable benefit for the design of our data analytics and application.

**Potential Risks**

The study will analyse your in-person responses to questions. Your responses will be recorded and some questions may be perceived to be intrusive or uncomfortable by you. The study may also require you to be untruthful or to conceal the truth, which may induce some level of discomfort for a short period of time. This discomfort is expected to be reversible upon stopping the experiment. You are allowed to stop the experiment and/or withdraw from the study at any time if it becomes uncomfortable with no penalty to you. Please see the section titled "Participation and Withdrawal" below for further information.

**Participation and Withdrawal**

Your participation in this study is entirely voluntary. You may withdraw from the study or terminate the study at any time at your own free will by informing the research staff. You may also refuse to answer any particular question that you do not wish to answer. Refusal to participate and withdrawal at any time during the study will not incur any penalties, however, monetary remuneration will not be provided to participants who withdraw from the study prior to completion. Checks on the automated interview system will be performed prior to your participation, however, the experiment may be terminated in the event of equipment failure. In such cases, reimbursement will be prorated based on the time spent during the experiment.

If you have changed your mind and no longer wish to participate, you can email the research staff to inform them of your decision to withdraw. Once we have received your email, someone from the research team will contact you to confirm your decision. Data from participants that withdraw from the study will be deleted and will not be used for further analysis.

**Confidentiality**

Any information that is obtained in connection with this study and personal data that can be used to identify you will remain confidential and will be disclosed only with your permission or as required by law. Your identifiable information and any data collected as part of the research will not be used or distributed, but de-identified information may be used for future research studies. Data collected will not be associated with your name. Confidentiality will be maintained by pseudonymisation: a randomly generated ID number will be used instead of your name when responses are recorded. These ID numbers will be kept in a secured storage medium and/or AWS' cloud computing infrastructure located in AWS' Singapore Data Centre, while data in hardcopy or physical form will be kept in secured premises. Besides the project team at AI Seer Pte. Ltd., only the Singapore Parkway Independent Ethics Committee (PIEC) will be granted direct access to your identifiable records for verification of research procedures and/or data, without violating your confidentiality, to the extent permitted by the applicable laws and regulations.

Data collected will be retained for data processing for a time period of no longer than 7 years. Reverse-identification of recorded responses will only be necessitated in the event of data loss, the need to extract further data, or requests for data to be withdrawn from the study. Any data identifying participants will not be shared with any third parties, published, or used for marketing purposes without consent. This study is sponsored by AI Seer Pte. Ltd., and data will be owned by AI Seer Pte. Ltd. solely for the purposes set out under the section titled "Purpose of the Study" above. By signing and dating this information and consent form, you consent to the collection, access, use, and disclosure of your information as described above.

**Whom can I contact about this study?**

If you have any questions or concerns about this study, you may contact the following staff:

Mr. Dennis Yap
CEO

Mobile number: +65 83050508

Ms. Veronica Sng
Lead Experimenter

Mobile number: +65 90608493

**Parkway Independent Ethics Committee (PIEC)**

An Institutional Review Board (IRB) or Ethics Committee (EC) is an independent group to protect the rights and well-being of subject volunteers participating in research. If you want an independent opinion of your rights as a research subject or to provide any feedback about this research, you may contact the Secretariat of Parkway Independent Ethics Committee (PIEC) at 6277 8272 or.

AI Seer Pte. Ltd.
UEN: 201942044R
24 St. Patrick's Road
Singapore 424146

ACKNOWLEDGEMENT, CONSENT AND ACCEPTANCE (PLEASE TICK):

- I have read and understood the contents of the Participant Information Sheet and this study consent form, and I have been given the opportunity to have my queries answered prior to the study.
- I fully consent to the terms of the study consent form and I agree to participate in this study on my own accord.
- I understand that I have the right to withdraw from this study at any point in time, and that I will not be required to explain my reasons for withdrawing.
- I understand that all the information I provide as part of this study will be treated in strict confidence and solely used for the purposes of research.

____________________________
Signature

____________________________
Name of participant

____________________________
Date

____________________________
Signature of staff

____________________________
Name of staff

____________________________
Date

AI Seer Pte. Ltd.
UEN: 201942044R
24 St. Patrick's Road
Singapore 424146

Version Date: 11th Sep 23
